# Synthetic bacteria with programmed cell targeting and protein injection suppress tumor growth *in vivo*

**DOI:** 10.1101/2024.04.22.590337

**Authors:** Alejandro Asensio-Calavia, Carmen Mañas, Alba Cabrera-Fisac, Eva Pico-Sánchez, Elena M. Seco, Starsha Kolodziej, Daniel S. Leventhal, José M. Lora, Beatriz Álvarez, Luis Ángel Fernández

**Author notes:** Department of Biosystems Science and Engineering, ETH Zurich, Schanzenstrasse 44, 4056 Basel, Switzerland. hC Bioscience, Inc., 645 Summer St., Boston, MA.

## Abstract

Bacterial living therapeutics (BLTs) hold promise for treating cancer and other human diseases because they can be engineered and transported into the microbiota (e.g., of tumors, gastrointestinal tract) to deliver therapeutic payloads. Current approaches rely on the natural tropism of the bacterial chassis used and trigger the local release of protein cargoes, typically through active extracellular secretion or bacterial lysis. BLTs capable of targeting specific cellular subsets and delivering payloads intracellularly might provide new therapeutic opportunities and improve efficacy while reducing off-target effects. We used synthetic biology to develop BLTs that can deliver defined cargo proteins into the cytoplasm of target cells. We designed a modular synthetic bacterium with programmed adhesion to cells by targeting defined cell surface antigen and armed with an inducible type III secretion system (T3SS) for injection of a protein cargo of interest. As a proof of principle, we programmed synthetic bacteria to recognize the epidermal growth factor receptor (EGFR) and inject the catalytic fragments of the potent adenosine diphosphate-ribosyltransferase toxins ExoA and TccC3. These BLTs demonstrated the ability to trigger robust tumor cell death *in vitro*. Intratumoral administration of these synthetic bacteria suppressed tumor growth *in vivo* and prolonged the survival of treated animals when the tumor cells were recognized by the engineered bacteria. These results demonstrate the potential of programming cell targeting and controlled protein injection for the development of effective and specific BLTs.

**Graphical Abstract:** 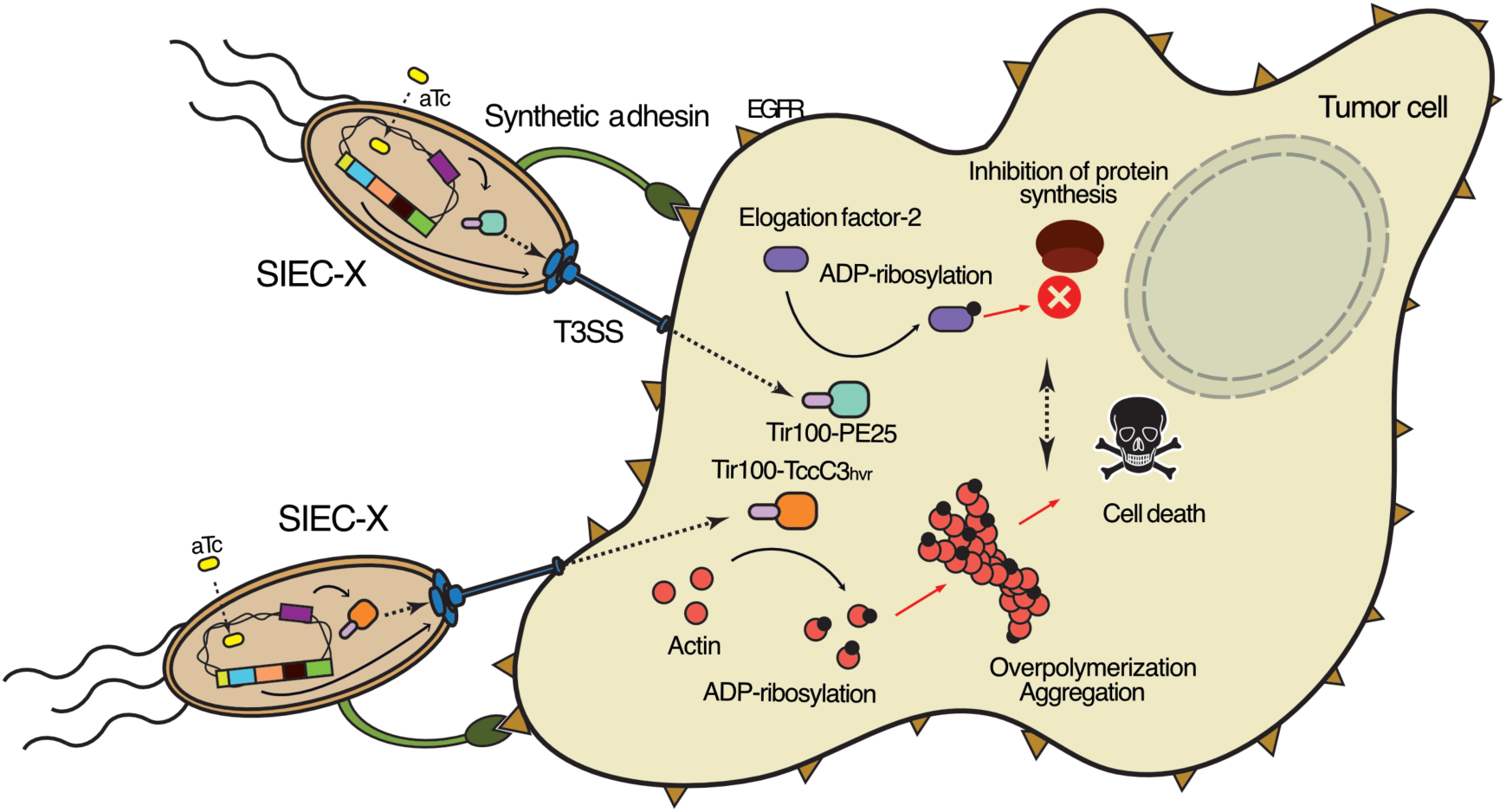

## Introduction

The development of living therapeutics (LTs) is a promising approach for treating multiple human diseases, including immunological and metabolic disorders, infections and cancer ^1, 2^. LTs are engineered cells that can produce therapeutic molecules such as antibodies, cytokines, metabolites, enzymes and cytotoxins against a specific disease ^3^. To enhance efficacy and minimize adverse effects, administered LTs should have the ability to control the production of the therapeutic payload. Contextual or temporal control of payload expression is usually accomplished with genetic circuits that respond to chemical or physical inducers, or the presence of disease-specific biomarkers ^4^. While control of the timing and location of payload production is key, it would also be advantageous to target specific cell subsets within a tissue. Chimeric antigen receptor (CAR) T-cell therapies are a notable example of LTs based on the reintroduction of T-cells engineered with selective tumor cell targeting specificity that trigger killing upon interaction of the CAR with its target on tumor cells ^5^.

Bacterial living therapeutics (BLTs) are an area of growing interest due to their relative ease of genetic manipulation and their potential for (re)introduction into the human microbiota ^6, 7^. Engineered bacteria can produce therapeutic proteins *in situ* that can combat viral and bacterial infections ^8, 9^, promote tissue repair ^10^, or reduce gut inflammation ^11–13^. Facultative anaerobic bacteria naturally colonize and proliferate in the hypoxic and immunosuppressive environment of solid tumors ^14^. This ability has spurred the engineering of bacteria for the local delivery of antitumor molecules ^15, 16^. Major recent achievements in this field have utilized synthetic biology to program bacteria for tumor immunotherapies, by depleting the levels of immunosuppressive metabolites ^17^, producing signaling molecules activating the immune response ^18^, releasing nanobodies (Nbs) that block antiphagocytic receptors and immunological checkpoints ^19^, or delivering synthetic antigens recognized by CAR-T cells ^20^. However, these successes invariably rely on the natural tropism of the bacterial chassis to colonize a specific niche (e.g., tumor or gut), where the therapeutic molecule is released locally into the environment either by active protein secretion or by bacterial lysis ^16^. Implementation of target cell recognition and an intracellular protein delivery system for therapeutic cargoes would increase the specificity and biosafety of BLTs, enabling selective targeting of specific cells in tumors and other diseases. Here, we have developed a BLT with programmed cell targeting and controlled injection of protein cargoes into tumor cells. To create this synthetic bacterium, we used a stepwise approach by stably integrating different genetic modules into the chromosome of a non-pathogenic *Escherichia coli* chassis to provide individual functionalities. These modules include protein delivery of cargo, regulation and cell targeting. The protein delivery module is based on injectisomes from the type III protein secretion system (T3SS) ^21, 22^. The injectisomes are large multi-protein complexes that act as nanosyringes in bacterial pathogens, delivering protein effectors into host cells during infection ^23^. Attenuated strains of pathogens with a natural T3SS have been used to deliver heterologous proteins into mammalian cells ^24–26^. Synthetic biology has enabled the expression of T3SS in non-pathogenic commensal *E. coli* strains ^27, 28^. We previously engineered the expression of T3SS injectisomes from enteropathogenic *E. coli* (EPEC) in *E. coli* K-12, generating the synthetic injector *E. coli* (SIEC) strain ^27^. SIEC encoded all structural proteins and chaperones necessary for assembling functional injectisomes in five integrated synthetic operons controlled by the isopropyl β-D-1-thiogalactopyranoside (IPTG)-inducible *tac* promoters (*Ptac*) ^27^.

In this work, we used SIEC as a bacterial chassis for the insertion of: i) a cargo module to translocate multiple heterologous proteins through the T3SS injectisome; ii) a regulatory module based on a three-repressor (3R) genetic switch ^29^ for the tight control of protein delivery *in vivo*; iii) a synthetic adhesin ^30^ module to enable selective attachment of the bacterium to tumor cells via binding to the human epidermal growth factor receptor (EGFR) ^31^. We have shown that this modular synthetic bacterium can deploy multiple heterologous proteins into the cytoplasm of tumor cells expressing EGFR. As a proof-of-principle, we demonstrate that this approach can be applied to the delivery of potent catalytic fragments of adenosine diphosphate (ADP)-rybosyltransferase (ART) toxins, causing an effective tumor cell death *in vitro* and a suppression of tumor growth *in vivo* when the synthetic bacterium recognizes tumor cells expressing EGFR.

## Results

### Engineering a cargo module for T3SS-dependent protein translocation

SIEC injectisomes were originally shown to translocate the natural Tir effector into HeLa cells^27^. More recently, we have shown that SIEC can translocate a protein fusion containing the N-terminal 20 amino acids of the EspF effector (EspF20), a T3S-signal, and the TEM-1 β-lactamase (Bla) lacking its natural Sec-dependent signal peptide ^32^. The enzymatic activity of Bla is commonly used as a reporter of protein translocation in mammalian cells by incubation with CCF2-AM, an esterified form of hydroxycoumarin linked to fluorescein by a β-lactam bond ^33^. Upon cell entry, mammalian esterases convert CCF2-AM to the negatively charged CCF2, which is retained in the cytosol and cleaved by the translocated Bla, switching its fluorescence emission from green (520 nm) to blue (450 nm). To develop an efficient translocation module of heterologous proteins in SIEC, we first compared the T3S-signal EspF20 with the N-terminal 30 and 100 residues of Tir, named Tir30 and Tir100, respectively. Since Tir100 contains the binding site for the multi-effector chaperone CesT ^34^, we placed the *cesT* gene in a bicistronic configuration with Tir100 in the construct (Fig. 1a). These T3S-signals were placed in-frame with an Nb (∼15 kDa) binding GFP, which was translocated by EPEC bacteria ^24^, and Bla (∼28 kDa) (Fig. 1a). A construct containing Bla and lacking the T3S-signal was also included as negative control. These protein fusions were expressed from multicopy pBAD plasmids (Table S2), with the L-arabinose (L-ara)-inducible P_BAD_ promoter, in the SIEC strain and its isogenic *ΔescN* mutant, which lacks the essential ATPase of the T3SS ^35^ (Table S1). The bacterial cultures were induced with IPTG (for expression of T3SS) and L-ara (for expression of Bla fusions) and used to infect HeLa cells *in vitro* (see Methods). The translocation levels of the Bla fusions in the infected cultures were determined by adding CCF2-AM and quantifying the ratio of fluorescence at 450 and 520 nm. These experiments showed that all T3S-signals in SIEC were functional for translocation of the Nb-Bla fusion in a T3SS-dependent manner, with the fusion having Tir100 signal and CesT exhibiting the highest levels of translocation (Fig. 1b). Western blot analysis of whole-cell bacterial protein extracts revealed similar expression levels of the Bla fusions in SIEC (Fig. S1). We also tested the SIEC translocation of the Tir100-Nb-Bla fusion in a different cell line. Mouse fibroblast 3T3 cells incubated with CCF2-AM showed blue fluorescence signals, characteristic of hydrolyzed CCF2, when infected with SIEC bacteria encoding the Tir100-Nb-Bla fusion and CesT (Fig. 1c). By contrast, 3T3 cells showed the green fluorescence of noncleaved CCF2 when infected with SIEC bacteria encoding Bla without the T3S-signal (Fig. 1c).

**Figure 1.**
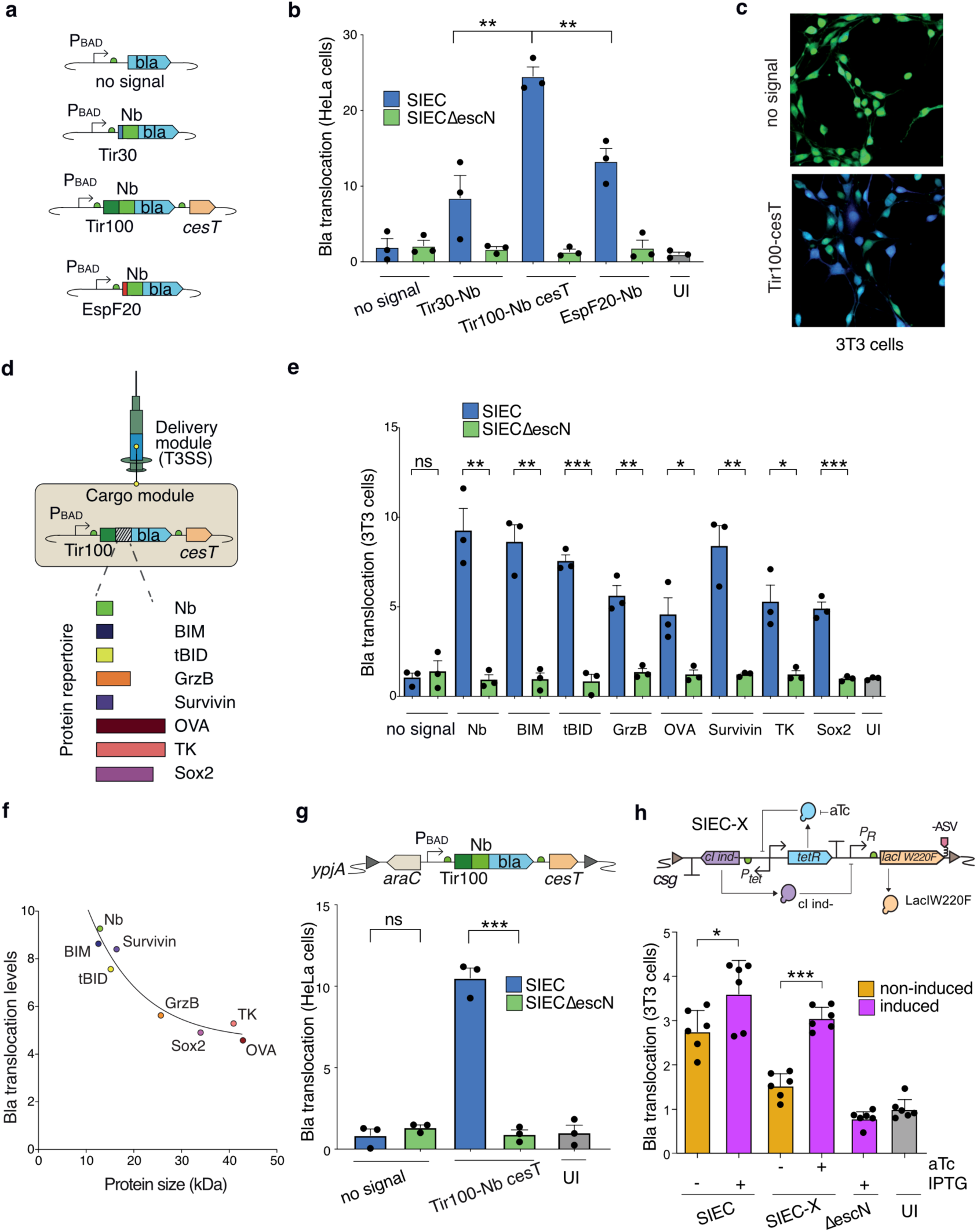
Engineering SIEC bacteria to deliver cargo proteins into the cytoplasm of mammalian cells. **a**, Schematic representation of the constructs in pBAD plasmids for the expression of different T3S signals (Tir30, Tir100, EspF20) fused to a nanobody (Nb) and β-lactamase (Bla), with the presence of the multicargo chaperone CesT with Tir100. **b**, **e, g, h**, Bla translocation levels were obtained by calculating the fluorescence emission ratio between 450 nm and 520 nm (blue emission/green emission). Each experimental condition was normalized by the emission ratio of uninfected cells (UI). Data are presented as the mean ± s.e. Unpaired two-tailed t-tests were used to evaluate differences between groups. (*) *P* <0.05, (**) *P* <0.01, (***) *P* <0.001, (****) *P* <0.0001. UI (uninfected control). **b**, Levels of Bla translocation in 3T3 cells infected with SIEC or isogenic *ΔescN* strains harboring Bla fusions to the indicated T3S signals in pBAD plasmid. Tir100-cesT versus Tir30 (*P*=0.0074) and versus EspF20 (*P*=0.0055) **c**, Fluorescence confocal microscopy of 3T3 mouse fibroblasts stained with CCF2-AM after 3 h of infection with SIEC bacteria carrying pBAD-bla (control) or pBAD-Tir100-Nb-bla, as indicated. **d**, Scheme representing the the cargo module Tir100-cargo-bla *cesT*, for delivery of heterologous proteins by SIEC, and the repertoire of heterologous proteins tested. **e**, Translocation levels of heterologous proteins. Each condition was compared with its correspondent *ΔescN* control: no signal (*P*=0.6098), Nb (*P*=0.0027), BIM (*P*=0.0016), tBID (*P*=0.0002), GrzB (*P*=0.0021), OVA (*P*=0.0251), Survivin (*P*=0.0032), TK (*P*=0.013), Sox2 (*P*=0.0005) **f**, Graph showing translocation levels of each protein in relation to its size (in kDa), with adjusted line corresponding to one-phase decay nonlinear fit. **g**, Translocation levels in HeLa cells of constructs Tir100-cargo-bla *cesT* and bla (no signal), integrated into the chromosomal *ypjA* locus of SIEC and isogenic *ΔescN* strains. Scheme of integrated construct *araC*-*P_BAD_*_Tir100-cargo-bla_*cesT* on top. Each condition was compared with its corresponding *ΔescN* control: no signal (*P*=0.3188), Tir100-Nb cesT (*P*=0.0001). **h**, Diagram on top represents parts and interactions of the regulation circuit 3R-X ^29^ integrated into the *csg* locus of SIEC to generate SIEC-X. Comparison of protein translocation levels in 3T3 cells by SIEC and SIEC-X strains, and isogenic *ΔescN* control, with overexpressed EspF20-Bla from pBAD plasmid (grown with L-ara prior to the infection) and their T3SS induced or not (+, -) with IPTG (SIEC) or aTc (SIEC-X) during infection. Each induced condition was compared to the non-induced: SIEC (*P*=0.0429), SIEC-X (*P*<0.0001).

We next investigated the delivery of a diverse repertoire of heterologous cargo proteins with different sizes and biological activities using pBAD vectors with Tir100 signaling and CesT. To do this, the Nb in the Tir100-Nb-Bla construct was replaced with DNA sequences encoding the following: the Bcl-2-interacting mediator of cell death (BIM) ^36^, the truncated BH3 interacting-domain death agonist (tBID) ^37^, granzyme B (GrzB) ^38^, ovoalbumin (OVA) ^39^, survivin ^40^, the herpes simplex virus thymidine kinase (TK) ^41^, and the mammalian transcription factor Sox2 ^42^ (Fig. 1d). SIEC and isogenic τ<<*escN* (T3SS defective negative control) strains carrying plasmids encoding these protein cargoes, Tir100-Nb-Bla (positive control), and Bla (negative control), were induced with IPTG and L-ara, as above, and used to infect 3T3 cells. Quantification of Bla activity in infected cells revealed a T3SS-dependent translocation of the heterologous cargo proteins at different levels (Fig. 1e), with an inverse correlation between their molecular mass and their level of translocation (Fig. 1f). Smaller cargoes of ∼12–20 kDa (e.g., Nb, BIM, tBID, survivin) showed a higher level of translocation than larger cargos of ∼35–45 kDa (e.g., OVA, TK, Sox2). These results demonstrated that the T3S signal Tir100, together with the co-expressed chaperone CesT, was able to promote the translocation of a variety of heterologous proteins.

As synthetic bacteria should ideally contain all genetic modules functioning stably in the chromosome, we designed the insertion of all genetic modules of SIEC in the chromosome using a markerless approach replacing nonessential genes in *E. coli* K-12 genome (Fig. S2) ^27^. We integrated the *araC*-P_BAD__Tir100-Nb-Bla_*cesT* genetic module into the *ypjA* locus of SIEC (Fig. S2) and SIEC*ΔescN* strains and tested Bla translocation from single copy expression after induction with IPTG and L-ara. The results showed translocation of the Tir100-Nb-Bla fusion in HeLa cells infected with SIEC bacteria but not with the isogenic *ΔescN* (Fig. 1g). We also detected the translocated polypeptide by western blot in cytoplasmic protein extracts from mammalian cells infected with SIEC with the integrated cargo module (Fig. S3). We then tested the integration into the *ypjA* locus of SIEC of genetic modules with *araC*-P_BAD_ encoding the natural EPEC effectors Tir, NleC, EspH, and Map fused to Bla and with downstream *cesT* ^43^. These effector-Bla fusions were also efficiently translocated by SIEC (Fig. S4), demonstrating the versatility of the integrated genetic module for the T3SS-dependent translocation of both natural effectors and heterologous protein cargos by SIEC bacteria.

### A regulatory module for tight control of protein delivery

Although the expression of protein cargoes could be controlled using the *araC*-P_BAD_ promoter, the expression of injectisomes in SIEC depends on Ptac promoters, which exhibit low levels of expression even in the absence of IPTG ^27^. In addition, IPTG has a short half-life *in vivo* and can be toxic ^44, 45^. To overcome these limitations, we have recently developed a genetic circuit based on three transcriptional repressors (TetR, cI and LacI^W220F^) ^29^ that allows more stringent regulation of Ptac promoters with anhydrotetracycline (aTc), an inducer suitable for *in vivo* use ^46, 47^. When this regulatory module, termed 3R-X, was integrated into the *csg* locus of the SIEC chromosome (Fig. S2), the resulting strain (termed SIEC-X) assembled injectisomes with reduced leakiness upon aTc induction ^29^. We therefore compared the levels of protein translocation in SIEC versus SIEC-X (Fig. 1h). To ensure high levels of protein cargo, the strains were transformed with a high-copy plasmid vector (pBAD_EspF20-Bla) and grown in medium with L-ara. Injectisomes were then induced (or not) by adding IPTG (for SIEC) or aTc (for SIEC-X) to the cultures (Fig. 1h). These experiments showed a significant protein translocation in SIEC in the absence of IPTG, with ∼70% of the level observed in the induced state. By contrast, SIEC-X showed a significant reduction in translocation of the protein cargo in the absence of inducer, whereas it showed similar levels of translocation to those of induced SIEC when aTc was added (Fig. 1h). We therefore implemented the 3R-X regulatory module in our synthetic bacterium to improve the control of injectisome expression *in vivo*.

### An adhesion module for attachment to target cells

We have previously reported synthetic adhesins (SAs) for *E. coli* based on the surface display of Nbs with specific antigen binding fused to the outer membrane (OM)-anchoring domain of intimin ^30^. *E. coli* bacteria constitutively expressing a SA binding GFP (SAgfp) attached selectively to HeLa cells expressing GFP on the plasma membrane (GFP-tm) ^30^. As the binding specificity of the SA can be switched by replacing the Nb, SAs are an appropriate adhesion module for programming cell targeting in our synthetic bacterium with T3SS, cargo and regulatory modules (Fig. 2a). EGFR is a receptor tyrosine kinase located on the cell membrane that is involved in cell proliferation and survival, among other functions^48^. As EGFR is a validated tumor-associated cell surface antigen, we used this receptor to develop targeted synthetic bacteria with T3SS-dependent protein delivery. We generated a SA binding EGFR (SAegfr) based on a previously selected Nb (Vegfr2) ^31^. SAegfr was placed under the constitutive promoter P_N25_ and integrated into the *flu* locus of SIEC-X (Fig. 2b; Fig. S2), and EcM1*lux*, a T3SS-deficient *E. coli* K-12 strain previously used for SA expression ^30^. Isogenic strains expressing a SA-binding Tir antigen (SAtir) were used as controls (Fig. 2b). We confirmed the expression and surface presentation of SAegfr in the bacterial strains by flow cytometry, which detected the C-terminal myc-tag (Fig. 2c). We then tested bacterial adhesion to cell lines with varying levels of expression for EGFR. EcM1*lux* expressing SAegfr adhered to 3T3 cells stably transfected with human EGFR (Her14 cell line) but not to non-transfected 3T3 cells (Fig. S5). Isogenic bacteria expressing SAtir did not bind to Her14 or 3T3 cells (Fig. S5). We tested the adhesion of SIEC-X strains expressing SAegfr and SAtir to Her14 and 3T3 cells, and to HCT116 human colon carcinoma cells. The expression of EGFR on the surface of these cell lines was compared by flow cytometry (Fig. 2d), which revealed higher levels in the transfected Her14 cells than in the HCT116 tumor cells. Nevertheless, after 1 hour of infection of these cell cultures, SIEC-X SAegfr bacteria adhered specifically and in high numbers to both EGFR+ cell lines (Her14 and HCT116) and not to the control 3T3 cell line (Fig. 2e). By contrast, SIEC-X SAtir bacteria failed to not attach to any of the cell lines tested (Fig. 2e). These results demonstrate that SAegfr is correctly expressed in SIEC-X and mediates specific bacterial adhesion to tumor cells expressing EGFR.

**Figure 2.**
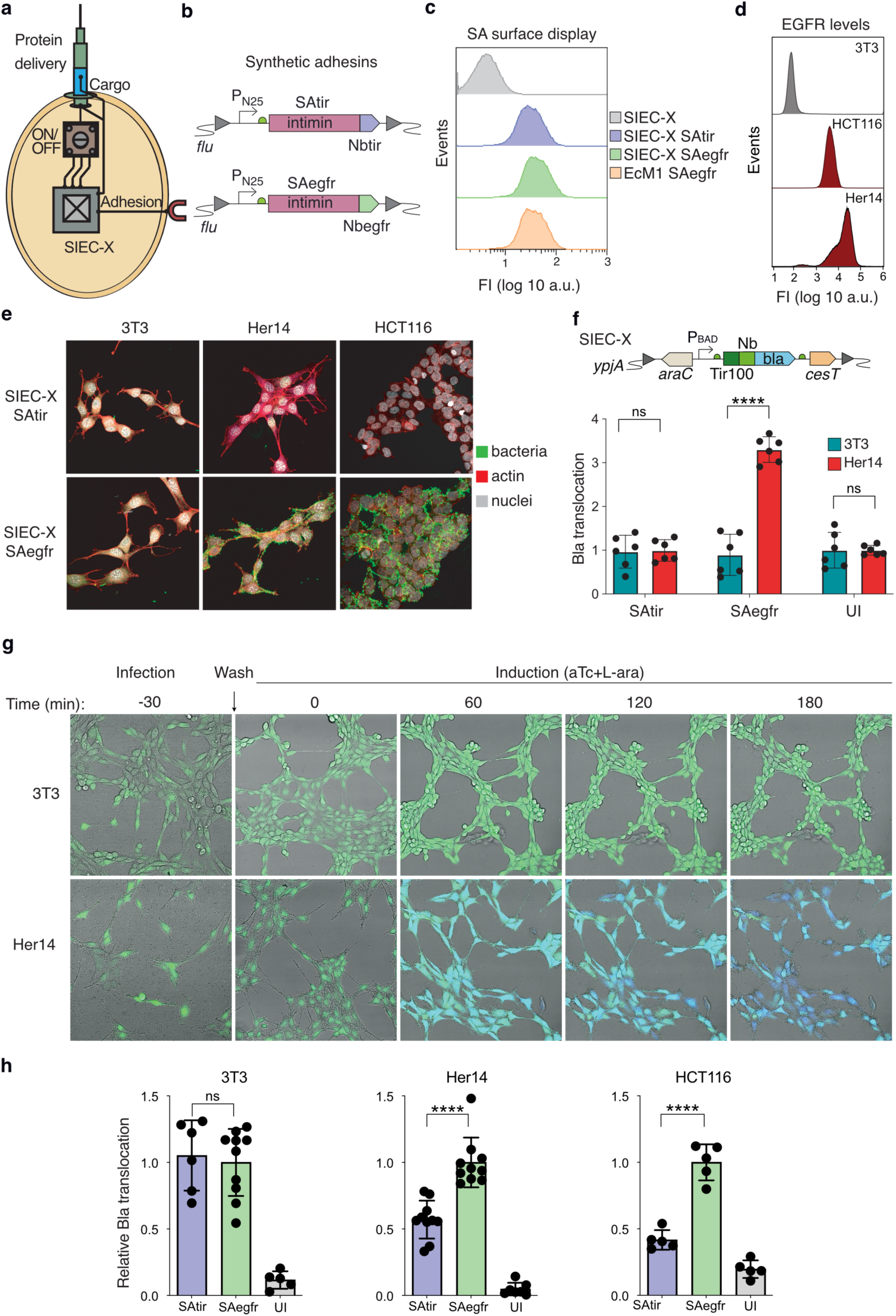
Synthetic adhesins in SIEC-X for cell-targeted translocation of protein cargos. **a**, Scheme of a synthetic bacterium with different functional modules for injection of protein cargos, regulation and specific adhesion to target cells. **b**, Synthetic adhesin constructs SAegfr, with Nb binding EGFR, and SAtir, with Nb binding tir (control). **c**, Surface display expression of SAegfr and SAtir in SIEC-X. Flow cytometry analysis of bacteria from the indicated strains stained with anti-myc monoclonal antibody (mAb) and secondary fluorophore-labeled anti-mouse antibodies (FI, fluoresce intensity; a.u., arbitrary units). **d**, EGFR expression in tumor cell lines 3T3 (negative), Her14, and HCT116. Flow cytometry analysis of cells labeled with anti-EGFR mAb and secondary fluorophore-labeled anti-mouse antibodies. **e**, Adhesion of SIEC-X bacteria with SAegfr or SAtir (control) to 3T3, Her14, and HCT116 cells. Fluorescence microscopy images after staining with anti-*E. coli* polyclonal antibodies (bacteria, green), phalloidin (actin, red), and DAPI (nuclei, grey). **f**, **h**, Bla translocation levels were obtained by calculating the fluorescence emission ratio between 450 nm and 520 nm (blue emission/green emission). Each experimental condition was normalized by the emission ratio of uninfected cells (UI) in **f**, or of the indicated cell line infected with SIEC-X carrying SAegfr in **h**. Data are presented as the mean ± s.e. Unpaired two-tailed t-tests were used to evaluate differences between groups. (*) *P*<0.05, (**) *P*<0.01, (***) P<0.001, (****) *P*<0.0001. **f**, Protein translocation of Tir100-Nb-bla fusion to Her14 cells expressing EGFR by SIEC-X bacteria with SAegfr. 3T3 (control) or Her14 (EGFR+) cells were infected with SIEC-X bacteria having the indicated SA (SAtir, SAegfr) and Tir100-Nb-bla fusion, washed to remove unbound bacteria and induced with aTc and L-ara. In each condition translocation to 3T3 was compared with the condition in Her14: SAtir (*P*=0.7777), SAegfr (*P*<0.0001). **g**, time lapse of the protein translocation process, showing fluorescence microscopy images of 3T3 and Her14 cells, preloaded with the ß-lactamase substrate (CCF2-AM), infected with SIEC-X SAegfr with Tir100-Nb-bla, washed, and induced with aTc and L-ara for the indicated time (in minutes). **h**, Enhanced protein translocation by SIEC-X with SAegfr to target cells expressing EGFR, Her14 and HCT116, and not to control 3T3 cells. The indicated cell line was infected with SIEC-X bacteria Tir100-Nb-bla, expressing SAegfr or SAtir, and induced with aTc and L-ara without a washing step to remove unbound bacteria. In each cell line condition SAtir was compared with SAegfr: 3T3 (*P*=0.7), Her14 (*P*<0.0001), HCT116 (*P*<0.0001).

We then investigated whether this specific bacterial attachment was translated into enhanced protein delivery to cells expressing EGFR. We infected 3T3 cells and Her14 cells with SIEC-X bacteria carrying the integrated cargo (*araC*-P_BAD__Tir100-Nb-Bla_*cesT*) and adhesion modules (SAegfr or SAtir). After 30 minutes of infection, unbound bacteria were washed off with fresh medium and the infected cultures were incubated for a further 3 hours in the presence of aTc and L-ara inducers. After this period, translocation of the Nb-Bla fusion was detectable only in Her14 cells infected with bacteria carrying SAegfr, and not in 3T3 cells or in infections of Her14 with SIEC-X bacteria carrying SAtir (Fig. 2f). We then performed real-time Bla translocation assays ^49^ in these cells using time-lapse fluorescence microscopy (Fig. 2g). To this end, Her14 and 3T3 cells were pre-loaded with the Bla (CCF2-AM) prior to infection. CCF2-loaded cells emit green fluorescence, which is converted to blue fluorescence upon Bla translocation. SIEC-X SAegfr bacteria efficiently attached to the surface of Her14 cells after 30 minutes of infection, and these cells emitted progressively higher blue fluorescence upon induction of injectisomes and cargo with aTc and L-ara, with cells showing strong blue fluorescence as early as 60 minutes after induction (Fig. 2g; Supplementary video S1). By contrast, SIEC-X SAegfr did not attach to 3T3 cells, and these cells retained the green fluorescence of uncleaved CCF2 throughout the experiment, indicating a lack of of translocation of the Bla fusion.

The above experiments indicated that the specific adhesion of SIEC-X SAegfr induces the preferential translocation of the cargo protein to cells expressing EGFR. However, a limitation in these *in vitro* assays is that the washing step removes most of the unbound bacteria. Therefore, we performed additional *in vitro* experiments to assess translocation levels to target cells under conditions where an identical number of the bacteria remained in the cell cultures throughout the infection. For this, 3T3, Her14 and HCT116 cells were infected with SIEC-X strains carrying SAegfr or SAtir and the cargo module *araC*-P_BAD__Tir100-Nb-Bla_*cesT*. Bacteria were allowed to infect these cell cultures for 60 minutes prior to the induction of injectisomes and cargo protein for a further 3 hours with aTc and L-ara, without any wash step during the process. Despite the stringent conditions of these *in vitro* infections, which resulted in a high bacterial density in all cell cultures during the induction period, the results showed that the presence of the specific adhesin SAegfr improved the translocation of the cargo protein to EGFR+ cells (Her14 and HCT116) by at least 2–3-fold compared with bacteria carrying the control SAtir (Fig. 2h). As expected, we also found that the level of protein translocation to 3T3 cells was not altered by the specificity of the SA expressed by the bacteria (Fig. 2h). Taken together, these findings indicate that SA provides a targeting mechanism via cell attachment and enhances the injection efficiency of the T3SS.

### Translocation of ADP-ribosyltransferase (ART) toxins induces tumor cell death

To determine whether the delivery of a specific protein cargo by SIEC-X could affect the biological activity of a target cell, we tested the induction of cell death upon translocation of bacterial ART toxins. These enzymatic toxins act in the cytoplasm by inactivating key proteins for cellular physiology through covalent modification with an ADP-ribose moiety^50^. Exotoxin A (ExoA) from *Pseudomonas aeruginosa* is a secreted ART toxin that internalizes into host cells and catalyzes the irreversible ADP-ribosylation of eukaryotic elongation factor 2 (eEF-2), inhibiting protein synthesis and, ultimately, inducing cell death ^51, 52^. TccC3 is another secreted ART toxin produced by *Photorhabdus luminescens* ^53, 54^ that internalizes into host cells and ADP-ribosylates F-actin, promoting actin polymerization, aggregation and condensation, which induces cell rounding and detachment from the substratum ^55^. The ART toxins contain distinct domains for cell internalization and for catalysis of ADP-ribosylation. The catalytic domains of both ExoA and TccC3 induce tumor cell death ^56, 57^. Notably, an immunotoxin containing a 38-kDa fragment of ExoA (PE38) including its catalytic domain is approved for clinical use ^58^. In the case of ExoA, the catalytic domain is located on the C-terminal ∼25-kDa fragment (PE25) (Fig. 3a), whereas in TccC3 the hypervariable region (hvr) is located on the C-terminal ∼30-kDa fragment (Fig. 3b). The gene segments encoding PE25 and TccC3_hvr_ were used to generate the genetic cassettes [*araC*-P_BAD__Tir100-PE25-Bla_*cesT*] and [*araC*-P_BAD__Tir100-TccC3_hvr_-Bla_*cesT*], which were integrated into the *ypjA* locus of SIEC-X SAegfr (Fig. S2). The translocation of these Bla fusions in HCT116 cells was compared with that of the Nb-Bla fusion by the corresponding SIEC-X SAegfr strain, which revealed that PE25 was translocated at similar levels to the Nb, whereas the TccC3 fusion was translocated at ∼1.5× higher levels (Fig. 3c). As expected, translocation of the toxin fragments was not detected in the isogenic *ΔescN* strains (Fig. 3c).

**Figure 3.**
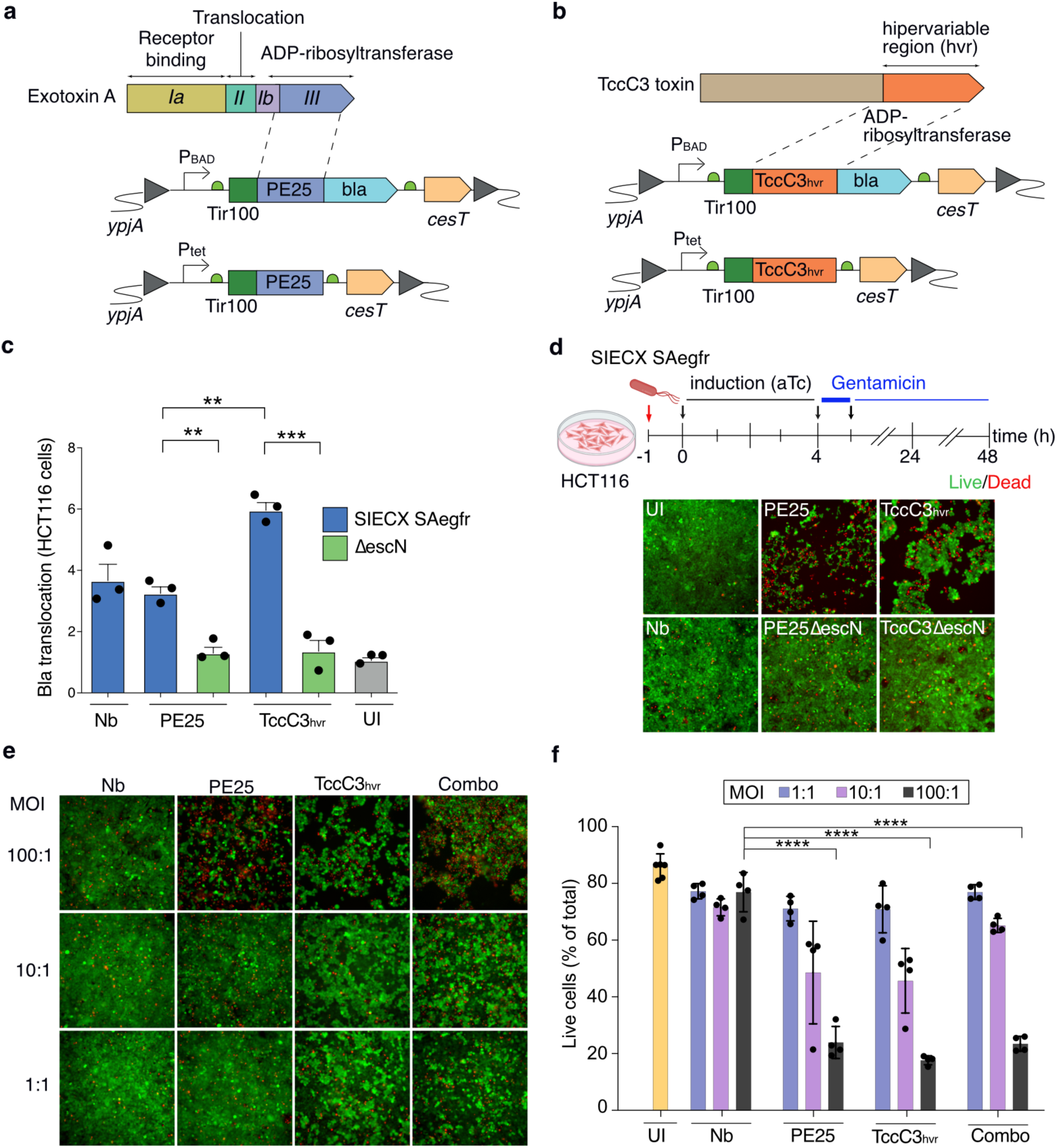
Translocation of ART-toxins by SIEC-X SAegfr induces tumor cell death. **a, b,** Schemes represent the domains of ExoA (**a**) and TccC3 (**b**) ART toxins, indicating the catalytic fragments inserted into the cargo module for SIEC-X translocation. **c**, **f**, Data are presented as the mean ± s.e. Unpaired two-tailed t-tests were used to evaluate differences between groups. (*) *P*<0.05, (**) *P*<0.01, (***) *P*<0.001, (****) *P*<0.0001 **c**, Bla translocation levels as in Figure 1. Translocation levels of the indicated toxin-Bla fusion to HCT116 cells, using Nb-Bla as positive control. SIECX SAegfr PE25 versus its ΔescN control (*P*=0.0025), TccC3_hvr_ versus its ΔescN control (*P*=0.0005), SIECX SAegfr PE25 versus TccC3_hvr_ (*P*=0.0013), **d**, Tumor cell death induced upon SIEC-X translocation of the indicated ART toxin fragment (PE25, TccC3_hvr_) or Nb as control. Scheme of the in vitro infection assay (top). Microscopy images of HCT116 cells stained for Live/Dead assay (Calcein-AM/Ethidium homodimer) 48 h post infection. Dead cells stain red whereas live cells stain green. Empty spaces within the monolayers indicate detached cells and/or reduced cell growth. Cell rounding is also visible among live cells with translocated ART toxins. **c, d,** infection with isogenic *ΔescN* strains are used as controls of the T3SS-dependent translocation of the toxins. **e,f**, effect of the multiplicity of infection (MOI) on the induction of tumor cell death. Infections with MOI 100:1, 10:1 and 1:1 (bacteria:cell) were carried out with SIEC-X SAegfr strains translocating Nb, PE25, TccC3_hvr,_ or with a 1:1 combination of bacteria translocating PE25 and TccC3_hvr_ (Combo). **e,** fluorescence microscopy images 48 h post infection of HCT116 stained for Live/Dead assay as in **d. f**, Graph shows percentage of live cells over total cells determined by flow cytometry analysis of HCT116 cells infected and stained for Live/Dead assay as in as in **e**. Nb versus PE25 (*P*<0.0001), versus TccC3 (*P*<0.0001) and versus Combo (*P*<0.0001).

We then evaluated the cytotoxic effect of the translocated PE25 and TccC3_hvr_ in HCT116 cells. We generated new integrative constructs lacking the Bla reporter and using the Ptet promoter instead of *araC*-P_BAD_ to drive expression of Tir100 fusions to Nb, PE25, and TccC3_hvr_, and downstream of CesT (Fig. 3a and 3b). Their integration into the *ypjA* locus of SIEC-X SAegfr allowed the induction of both T3SS injectisomes and cargo proteins with aTc as a single inducer. These bacterial strains, and isogenic *ΔescN* mutants, were used to infect HCT116 cells *in vitro*. After 1 hour of infection to allow adhesion, remaining unbound bacteria were washed from the cultures, and fresh medium containing aTc was added for a further 4 hours to enable payload expression and injection. After this period, some samples were evaluated by fluorescence microscopy using a Live/Dead assay in which live cells were stained with Calcein-AM (green) and dead cells with ethidium homodimer-1 (red). Parallel unstained samples were further incubated in culture medium with gentamicin (100 μg/mL) for 1 hour to kill bacteria, followed by further incubation in culture medium with a lower concentration of gentamicin (10 μg/mL) to prevent bacterial growth. At 24 and 48 hours after the initial infection, the cell samples were also stained for Live/Dead cells and examined by fluorescence microscopy (Fig. 3d and Fig. S6). These assays revealed an evident cytotoxic effect of both translocated ART toxins in HCT116 cells, with a clear cell rounding and cell detachment phenotype of cultures infected with bacteria translocating TccC3_hvr_, and a large proportion of dead cells in cultures infected with bacteria translocating PE25. Cell-rounding phenotype induced by TccC3_hvr_ was likely due to actin-crosslinking provoked by its ADP-ribosylating activity. Cell rounding was observed as early as 4 hours after induction of bacteria translocating TccC3_hvr_ and increased in samples incubated for 24 and 48 hours (Fig. S6 and Fig. 3d). Translocation of PE25 caused massive cell death, likely after inhibition of protein translation, which is known to trigger apoptosis by several mechanisms ^51^. Dead cells were not detectable after 4 hours of infection with PE25-translocating bacteria but were abundant after 24 hours and prevalent after 48 hours (Fig. S6 and Fig. 3d). At these late times after infection, cell rounding was also observed in the remaining live cells of these cultures. In contrast, cells infected with isogenic *ΔescN* mutants, or with bacteria translocating the Nb, showed a normal proliferation of live cells, as in uninfected (UI) controls, and no cytopathic phenotypes (Fig. 3d). These results demonstrate the biological activity of translocated ART toxins in the cytosol of HCT116 tumor cells. The cell rounding phenotype induced by TccC3_hvr_ was likely due to actin-crosslinking provoked by its ADP-ribosylating activity.

To investigate the dose-response effect of the translocated toxins, we performed Live/Dead assays on HCT116 cells using different multiplicities of infection (MOIs), from the originally used 100:1 to lower doses of 10:1 and 1:1. In addition to the previous experimental groups (i.e., UI, Nb, PE25 and TccC3_hvr_), we introduced a new group by infection with a combination of equal numbers of bacteria translocating PE25 and TccC3_hvr_ (Combo). Cell cultures were infected for 4 hours, treated with gentamicin as above, and analyzed after 48 hours by fluorescence microscopy (Fig. 3e) and by flow cytometry, to quantify live and dead cells (Fig. 3f). As expected, cytotoxicity and the percentage of dead cells increased with the bacterial dose for each individual toxin as well as for the Combo, being maximal at the highest MOI (100:1). Similar to the 4-hour infection results, the cell-rounding phenotype induced by TccC3_hvr_ was also evident at low MOIs (10:1 and 1:1) (Fig. 3e). The phenotype of HCT116 cells infected with the Combo was a combination of the cell death and cell rounding phenotypes of both toxins, with the latter detectable at low MOI (1:1) (Fig. 3e). Remarkably, despite the clear phenotypic differences observed by fluorescence microscopy, cultures quantified by flow cytometry at 48 hours showed a similar percentage of dead cells after infection with bacteria delivering PE25 or TccC3_hvr_ (Fig. 3f), suggesting that flow cytometry might more sensitive than the microscopy at detecting dead cells. Previous studies have also reported that TccC3 and its catalytic domain induce cell death by “freezing” the actin cytoskeleton and actin clustering, ultimately activating several cell death mechanisms ^57,59^. No significant levels of cell death were found by flow cytometry in the UI controls, nor in tumor cells infected with bacteria translocating the Nb or infected with a low MOI of TccC3_hvr_-translocating bacteria (Fig. 3f).

### Cell-targeted bacteria injecting ART toxins suppress tumor growth *in vivo*

To evaluate the anti-tumor capacity of SIEC-X strains with ART toxins *in vivo*, we used a human tumor xenograft model based on the subcutaneous implantation of HCT116 cells (3×10^6^) into athymic Nude mice (Fig. 4a). Tumor-bearing mice were randomly divided into different experimental groups (n=10) to receive PBS (control), SIEC-X SAegfr bacteria carrying the cargo proteins Nb, PE25, TccC3_hvr_, and a combination of bacteria translocating PE25 and TccC3_hvr_ (Combo). All mice were treated intratumorally (i.t.) on days 0, 3 and 6, at a dose of 10^9^ CFU/mouse (Fig. 4a). The Combo condition contained 5×10^8^ CFU of each SIEC-X injecting toxin. For induction of T3SS and cargo, aTc was administered daily by intraperitoneal (i.p.) injection. Tumor volumes were monitored for 15 days after the first infection (d.p.i.), when the first deaths occurred in the PBS and Nb groups. Notably, at 15 d.p.i. we found significant differences in tumor volume between the combo treated mice and all other experimental groups (Fig. 4b). The calculated tumor growth rate was ∼10 mm^3^/day for the Combo group compared with a tumor growth rate of ∼25 mm^3^/day for the other groups. However, after ∼10 days from the last bacterial dose (i.e., 16 d.p.i.) the tumor growth rate increased in the Combo group (Fig. S7). We also found that the probability of survival was higher in the Combo group than in the other groups up to 30 days post d.p.i. (Fig. 4c).

**Figure 4.**
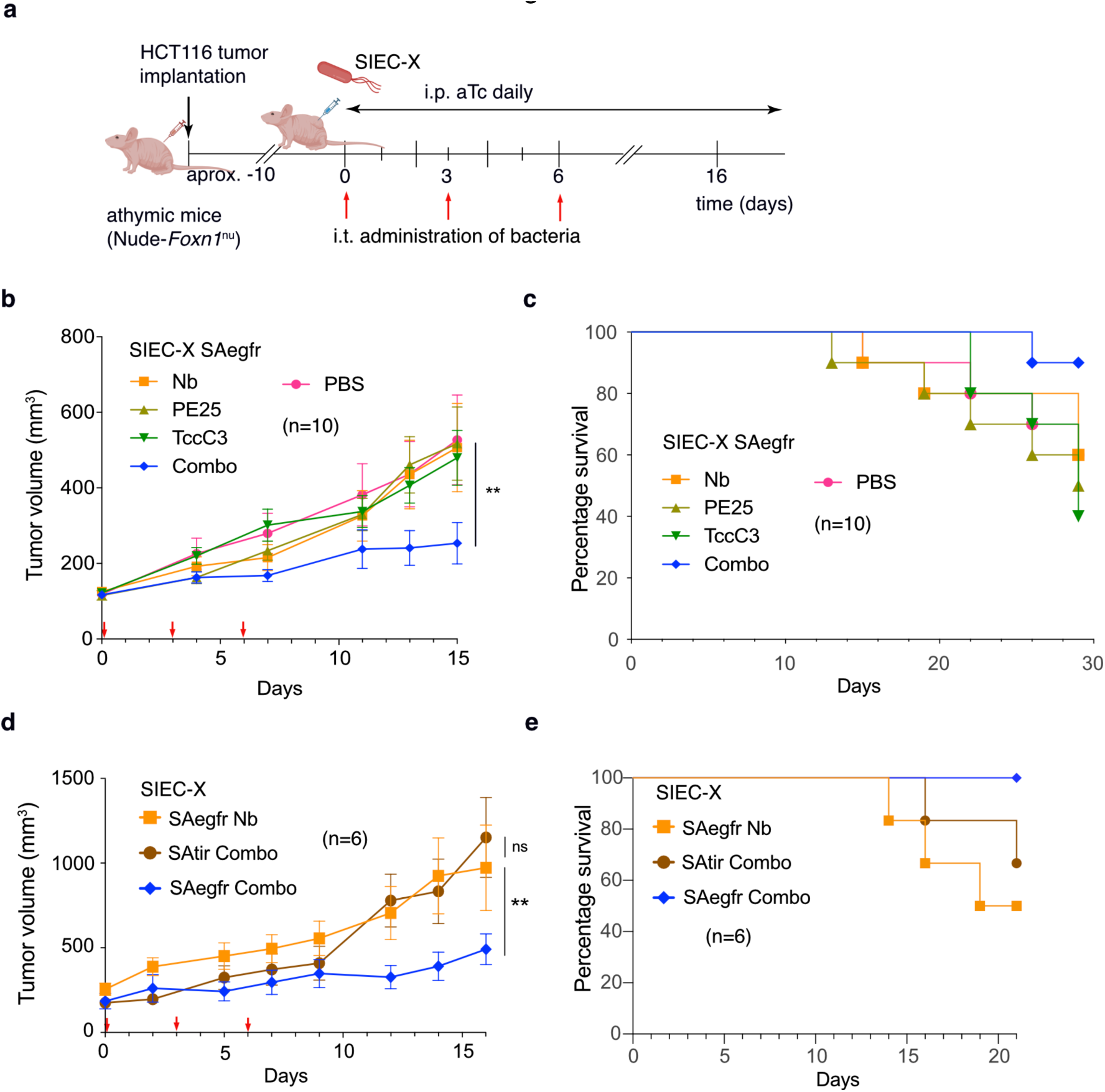
SIEC-X SAegfr bacteria translocating both ART toxins reduce tumor growth *in vivo*. **a**, Scheme overview of the in vivo experiments. Athymic Nude mice were subcutaneously implanted with HCT116 colon tumor cells 10–15 days prior to the intratumoral (i.t.) administration of the indicated SIEC-X bacteria (days 0, 3, 6). Bacterial doses: 10^9^ CFU/mouse in **b, c**; 10^8^ CFU/mouse in **d, e**. From the day of the first bacterial inoculation, mice were administered daily with aTc intraperitoneally (i.p.) to induce T3SS and protein cargo. **b, d,** Data are presented as the mean ± s.e of **b**, n=10 mice per group or **d**, n=6 mice per group. Statistical significance between groups determined by two-way ANOVA. (*) *P*<0.05, (**) *P*<0.01, (***) *P*<0.001, (****) *P*<0.0001 **b**, Graph showing 15-day evolution of tumor volume after bacterial (or PBS) administration, with lower red arrows indicating administration days. PBS versus Combo (*P*<0.01), **c**, percentage of survival mice from each group throughout experiment. **d**, relevance of the specific adhesion to tumor cells for tumor growth reduction, showed by 16-day evolution of tumor volume after the administration of the different strains. SAegfr Nb versus SAtir Combo (*P*=0.4946), SAtir Combo versus SAegfr Combo (*P*=0.0012) **e**, survival throughout the experiment. **c, e**, Statistical significance between groups inferred by Log-rank Mantel-Cox test. **c**, Nb versus Combo (*P*=0.1364). SAegfr Nb versus SAegfr Combo (*P*=0.0554).

Since the adhesion of SIEC-X bacteria to target cells enhances the translocation of protein cargos *in vitro* (Fig. 2h), we investigated whether the specific targeting of HCT116 tumor cells by SAegfr would also have a positive effect on the observed reduction in tumor growth induced by the Combo group *in vivo*. To this end, we designed an *in vivo* experiment as described above comparing Combo groups carrying SAegfr and SAtir (binding null control). We constructed isogenic SIEC-X strains translocating PE25 or TccC3_hvr_ cargos in which SAegfr was replaced by the control adhesin SAtir. These bacterial strains were used in a combination called SAtir Combo. After subcutaneous implantation of HCT116 cells, tumor-bearing mice were randomized into three groups (n=6) receiving i.t. bacteria of the Combo SAegfr, Combo SAtir, or SIEC-X SAegfr translocating Nb (payload null control). To evaluate efficacy at a lower bacterial dose, in this experiment bacteria were administered at 10^8^ CFU/mouse on days 0, 3, and 6. As before, the aTc inducer was administered i.p. daily to all mice and tumor volume was monitored. Despite the 10-fold lower bacterial dose used in this experiment, we found an effective reduction in tumor growth in the Combo SAegfr group at 16 d.p.i. (Fig. 4d and Fig. S8). At this time point, the Combo SAtir group did not show a significant reduction in tumor growth compared with the SAegfr Nb control group (Fig. 4d). Survival rate was also higher in the Combo SAegfr group than in the Combo SAtir and control groups (Fig. 4e). Harvested tumors from animals of all experimental groups contained similar numbers of viable bacteria (∼10^9^ CFU/g of tumor) irrespective of the strain administered to the mice (Fig. S8). These results demonstrated that effective SIEC-X anti-tumor activity *in vivo* is dependent on both targeting/adherence to tumors cells (via SAegfr) and delivery of a cytotoxic payload (via T3SS-depedent injection of PE25 and TccC3_hvr_ toxins).

## Discussion

Targeting tumor cells by recognizing a cell-surface antigen and generating a cytotoxic response are the key elements of CAR-T cells ^5^ for achieving durable therapeutic responses in hematological malignancies. Major efforts are underway to improve CAR-T cell therapies for the treatment of solid tumors, including their combination with engineered bacteria ^20^. The SIEC-X bacteria reported here provide a novel therapeutic approach in this global effort against solid tumors and represent a step forward in BLTs.

Several studies have reported the anti-tumor effects of bacteria engineered to release (by active secretion or lysis) a wide range of therapeutic molecules into the tumor microenvironment, including cytolysins ^60^, cytokines ^61^, apoptosis-inducing factors ^62^, prodrug-converting enzymes ^63^, Nbs ^19^, and immunostimulatory signaling molecules ^18^. These approaches rely on the natural tropism and proliferation of bacteria in solid tumors ^14^, and the consequent accumulation of the therapeutic molecules in the tumor microenvironment. For intracellular bacteria, such as *Salmonella*, both its natural tumor tropism and cell invasion capabilities have been used to deliver intracellular protein payloads into tumor cells ^64^. However, this approach lacks selective invasion of a target cell, and protein delivery is achieved by lysis of the invading bacteria, releasing all bacterial contents within the cell. By contrast, SIEC-X bacteria are selective in their recognition of the target cell through the binding of a cell-surface antigen and in the delivery of the protein payload, using the T3SS to inject specific cargo proteins.

The T3SS is used by bacterial pathogens to deliver protein effectors into host cells, thereby subverting cellular functions in favor of infection ^23^. Pathogenic strains were initially used to deliver heterologous proteins into the cytoplasm of mammalian cells (for review ^26^). However, even attenuated strains of pathogens pose a biosafety risk and compromise the specificity of cargo delivery as they simultaneously inject natural T3SS effectors. This limitation can be partially overcome by generating effector-less strains ^65, 66^, but the risk would still remain that uncharacterized effectors might be present in their genomes. Therefore, a more rational approach to increase the safety and specificity of protein delivery is to engineer the expression of T3SS in a non-pathogenic bacterial chassis that delivers only a defined therapeutic protein cargo. To this end, we developed the SIEC strain, which integrates five operons encoding EPEC injectisomes into the chromosome of *E. coli* K-12 and demonstrates translocation of the Tir effector ^27^. In a parallel study, a plasmid-based system encoding the T3SS of *Shigella flexneri* was developed in *E. coli* K-12 and several human transcription factors were translocated ^28^. This *E. coli* strain was also used for the T3-dependent extracellular secretion of Nbs in the gut ^12^.

In this work, we have extended the use of synthetic bacteria with engineered T3SS for the intracellular delivery of cargo proteins into target cells and demonstrated their potential *in vitro* and *in vivo*. Translocation of various heterologous proteins by SIEC was achieved by generating a bicistronic cargo module with the T3S signal Tir100 fused to the cargo sequence and followed by the CesT chaperone. Initially we used this module to translocate a Nb, several pro-apoptotic factors (BIM, tBID, and Grz-B), two antigenic proteins (OVA, Survivin), a pro-drug converting enzyme (HSV-TK), and a human transcription factor (Sox2). An inverse correlation between cargo size and translocation efficiency was observed, as small protein cargos (<20 kDa) were translocated with high efficiency (e.g., Nb, BIM, tBID, Survivin). It was also interesting to demonstrate that this cargo module was effective when integrated in a single copy into the chromosome of SIEC, as shown for the translocation of the Nb, the effectors Tir, NleC, EspH, and Map, and the two ART toxin fragments. This versatility of the cargo module is an important aspect in the development of SIEC strains for different applications.

Protein translocation was controlled in SIEC by incorporating into the chromosome a regulatory module that responds to aTc, an inducer that works both *in vitro* and *in vivo* ^18,47^. Insertion of the 3R-X circuit reduces the leaky expression of injectisome components in SIEC ^29^. We show here that SIEC-X also provides improved control of protein delivery to mammalian cells, significantly reducing protein translocation in the absence of the T3SS inducer. Notably, the tighter control of injectisomes in SIEC-X did not prevent rapid (∼30 minute) and efficient protein translocation when aTc was added, as shown by real-time Bla translocation assays.

Another essential element is the adhesion module in SIEC-X, which controls the specific attachment of the synthetic bacteria to the target cells. The adhesion module is based on SAs with an exposed Nb domain that binds to a cell surface antigen ^30^. An SA with a Nb binding GFP induced the specific adhesion of *E. coli* K-12 bacteria to HeLa cells expressing GFP on their surface and improved the colonization of tumors derived from these cells in athymic Nude mice ^30^. Here, we generated an SA binding EGFR (SAegfr), a cell surface antigen expressed by human tumors of epithelial origin ^67^, and demonstrated that this adhesion module can be implemented in SIEC-X to drive bacterial attachment to tumor cells expressing EGFR. Furthermore, SIEC-X bacteria expressing SAegfr elicited efficient cell-targeted protein translocation through a combination of binding specificity and enhanced translocation to EGFR+ cells. EGFR is currently being targeted with antibodies and small inhibitors in therapies for metastatic colon cancer and head and neck cancer ^68, 69^. The clinical validation of EGFR led us to use it as proof-of-principle with SIEC-X bacteria, but SAs that bind other tumor-associated cell surface proteins can be constructed using Nb domains of the desired specificity ^70^. In addition, SAs with different target specificities could be combined in SIEC-X, in a single bacterium or as a combination of strains, to bind different tumor cell surface antigens, thus reducing the possibility of tumor antigen escape mutants ^71^.

We have shown that targeted SIEC-X bacteria can deliver the catalytic fragments PE25 and TccC3_hvr_ ART toxins into HCT116 colon cancer cells in a T3SS-dependent manner. In their native form, the ExoA and TccC3 toxins are secreted by *P. aeruginosa* and *P. luminescens*, respectively, and are internalized into host cells to ADP-ribosylate specific protein targets, eEF-2 ^51, 52^ and F-actin ^53, 54^, respectively. Delivery of these toxins by SIEC-X with SAegfr resulted in effective HCT116 tumor cell death *in vitro*, reduced tumor growth *in vivo* and increased survival of the treated animals. The concept of targeted delivery of cytotoxins to cancer cells has been extensively explored in the past with immunotoxins ^72^. High clinical success in the treatment of treat hairy cell leukemia has been achieved with an immunotoxin containing a 38-kDa catalytic fragment of ExoA (PE38) fused to an anti-CD22 antibody ^58, 73^. ExoA-PE38 inhibits protein translation by ADP-ribosylation of eEF-2, which induces apoptotic cell death ^51, 52^. This encouraged us to use PE25 as a cargo protein in SIEC-X. We also attempted to trigger cell death by a different mechanism using the catalytic domain of TccC3_hvr_, which ADP-ribosylates F-actin, causing cell rounding and, ultimately, cell death ^57, 59^. Analysis of *in vitro* cultures infected with PE25-delivering bacteria showed marked cell death, in a dose-dependent manner after 24 h.p.i. By contrast, bacteria delivering TccC3_hvr_ predominantly induced a cell rounding and cell detachment phenotype as early as 4 h.p.i. and at low MOI (1:1 to 10:1) after 48 hours. This higher activity of TccC3_hvr_ could be related to its higher translocation compared with PE25, as shown with Bla-fusions, or to their different mechanisms of action. Despite these differences, both ART toxin domains showed similar levels of cell death in flow cytometry analysis, suggesting that cells intoxicated with TccC3_hvr_ also become highly sensitive and are induced to cell death.

We have demonstrated in a xenograft model in athymic mice with subcutaneous tumors of HCT116 cells that i.t. administration of a combination of two SIEC-X SAegfr strains, each translocating the individual toxin fragments PE25 and TccC3_hvr_, results in a reduction in tumor growth *in vivo* and increased survival rates in mice treated with this combination. Compared with control groups, no reduction in tumor growth was observed when PE25 or TccC3_hvr_ were individually administered. Thus, the cell death generated by each toxin alone was not sufficient to suppress tumor growth in this *in vivo* model, highlighting the importance of targeting tumors using multiple independent cytotoxic mechanisms. The synergistic activity of the ART toxins observed *in vivo* contrasts with the *in vitro* results, where the individual translocation of either toxin fragment resulted in efficient cell killing, and their combination by co-infection did not result in higher levels of cell death *in vitro*. It is likely that impacts of TccC3_hvr_ (and synergy with PE25) may be more pronounced *in vivo* where disruption of cytoskeleton could have a larger impact on cell viability ^74, 75^.

Remarkably, the *in vivo* experiments demonstrated that the efficacy of the SIEC-X bacteria expressing ART toxins was only achieved when the bacteria expressed the SAegfr, which attaches to the tumor cell, and not when a control SA (SAtir) was expressed. In the absence of tumor cell attachment, the translocation levels by SIEC-X bacteria appeared insufficient to suppress tumor growth at the macroscopic level in this model. Therefore, the targeting of tumor cells by SAegfr is critical to achieve the antitumor effects *in vivo*. This is consistent with the *in vitro* results, where the expression of SAegfr in SIEC-X increased the adhesion and translocation efficiency of the protein cargo in EGFR+ cells (Her14, HCT116). We believe that targeting cells with SAs may result in more effective delivery of the cytotoxic molecules *in vivo*, where bacteria are in close contact with the tumor cells. Overall, these results demonstrate the efficacy of an anti-tumor treatment based on cell-targeted synthetic bacteria injecting a combination of ART toxins with a T3SS, providing proof-of-principle for cell-targeted BLTs that could be developed against cancer and other human diseases.

## Methods

Detailed description of the methods used can be found in the Supporting Information file, including Tables of bacterial strains (Table S1), plasmids (Table S2), oligonucleotides (Table S3), antibodies and fluorophores (Table S4), and Supporting References.

### Statistical analysis

The statistical analysis in this work were performed using Prism software versions 9 and 10 (GraphPad Software Inc.). The specific statistical test used for each experiment and *P* values are detailed in the corresponding figure legends.

### Ethic statement

Experiments with mice performed in the CNB-CSIC Animal House Facility (ref. ES280790000182) followed the protocols approved by the Ethics Committee for Animal Experimentation of CSIC and authorized by the Division of Animal Protection of the Comunidad de Madrid (project reference PROEX 074/18). Animals were handled in strict accordance with the guidelines of the European Community 86/609/CEE. The experiments performed at Synlogic were reviewed and approved by Mispro’s Institutional Animal Care and Use Committee (Mispro Biotech Services, 400 Technology Square, Cambridge, MA, 02139), in compliance with Animal Welfare Act of US and EU legislation for experimental animals’ procedures.

### Supporting Figures and video

Supporting Figures S1 to S8 and associated Figure Legends can be found in the Supporting information file. Supporting video S1 was generated with frames from a real-time β-lactamase translocation assay of SIEC-X SAegfr Tir100-Nb-Bla bacteria infecting 3T3 and Her14 cells.

### Data and materials availability

All the data generated in the study are included in the present manuscript and the associated Supporting Information file. At the time of acceptance of this manuscript for publication, all raw data used to generate the final Figures will be available from a public repository (https://github.com/). DNA sequences of genetic constructs will be available from Genbank (https://www.ncbi.nlm.nih.gov/genbank/). All the materials described are available from the corresponding author upon reasonable request for research purposes.

## Acknowledgements

We thank Carlos Piñero-Lambea, David Ruano-Gallego and Gad Frankel for scientific discussions. We also thank Ning Li for technical discussions on strain engineering The excellent technical support of CNB-CSIC core scientific facilities “Advanced Light Microscopy” and “Flow Cytometry” is greatly appreciated. We thank Dr Kenneth McCreath editing the manuscript.

## Authors’ contributions

L.A.F. conceived the study and secured funding. A.A.C., C.M., A.C.F., E.P.S., E.M.S., and S.K., performed the experiments. All authors designed the experiments, analyzed the results, and interpreted the data. B.A. and L.A.F. supervised results and reproducibility. A.A.C. and B.A. wrote the initial draft manuscript and figures. LAF prepared the final manuscript and figures. All the authors revised and approved the final manuscript.

