## Supplemental Information for "Synthetic bacteria with programmed cell targeting and protein injection suppress tumor growth *in vivo*"

* Corresponding author:

- Supporting Methods
- Table S1. Bacterial strains.
- Table S2. Plasmids.
- Table S3. Oligonucleotides.
- Table S4. Antibodies and fluorophores.
- Supporting References.
- Supporting Figures S1 to S8.

### Supporting Methods

**Bacterial culture conditions**. The *E. coli* strains used in this work are listed in Table S1. Bacteria were grown in Lysogeny Broth (LB) [1] unless otherwise specified. For solid media formulations, LB agar plates containing 1.5% w/v agar were used. For liquid cultures, bacteria were incubated at 37ºC with shaking, as specified. When selection was needed, antibiotics were added at the following concentrations: kanamycin (Km) at 50 µg/mL, spectinomycin (Sp) at 50 µg/mL and ampicillin (Amp) at 150 µg/mL; all were purchased from Duchefa-Biochemie. To kill bacteria in *in vitro* infections, gentamicin (Gm; Gibco-Thermo Fisher Scientific) was used at 100 µg/mL for 1 hour followed by 10 µg/mL for 24 or 48 hours, as indicated. Chemical inducers used were as follows: L-arabinose (L-ara; Sigma) at 0.4% (w/v); isopropyl β-D-1-thiogalactopyranoside (IPTG; Sigma) at 0.1 mM; anhydrotetracycline (aTc, TOKU-E) at 200 ng/mL.

**Plasmid constructs and *E. coli* chromosomal engineering.** Plasmids used in this work are listed in Table S2. Visual glyphs of the biological design representations were created following the Synthetic Biology Open Language standards guidelines (SBOL Visual) [2]. Plasmids were constructed using Type II restriction enzyme cloning [3]. Oligonucleotides used as primers were purchased from Sigma-Aldrich and are listed in Table S3. Amplification of inserts was carried out by PCR using the proofreading DNA polymerase Herculase II Fusion (Agilent Technologies), followed by electrophoretic DNA band isolation. Ligation of plasmid backbones and inserts was performed overnight using T4 DNA ligase (Roche). All generated constructs were first screened for the presence of the insert by PCR amplification using NZYProof 2x Green Master Mix (NZYtech) and were sequenced using the Sanger chain-terminator method (Macrogen) with specific primers (see sequencing primers in Table S3). *E. coli* DH10B-T1R [4] was used as cloning strain for replicative plasmids such as pBAD18 and derivatives. For cloning and propagation of suicide pGE-plasmid derivatives containing the conditional pir-dependent R6K origin of replication [5], the *E. coli* strain BW25141 [6] was employed. Non-suicide plasmids were introduced into *E. coli* with Transformation and Storage Solution (TSS) [7] or by electroporation of electrocompetent cells [3]. Transformation of *E. coli* with suicide plasmids for chromosomal integration was always performed by electroporation. Site-specific deletions and insertions in the chromosome of *E. coli* were performed based on expression of *I-SceI* endonuclease [8] with a marker-less strategy of genome edition, as previously described [9, 10]. After the integration process, individual Km-sensitive colonies were screened by PCR with specific oligonucleotides to identify those with the desired modification in their chromosome (i.e., deletion, insertion, substitution). If necessary, bacterial chromosomes were isolated and the integrated region amplified with Herculase II Fusion, followed by agarose gel purification of the corresponding amplicon for Sanger sequencing with specific primers. The *E. coli* strains engineered for this work are listed in Table S1.

**Western Blot.** Sodium Dodecyl Sulfate-Polyacrylamide gel electrophoresis (SDS-PAGE) was carried out following standard methods on the Miniprotean III system (Bio-Rad) [3]. Proteins were boiled 10 minutes in reducing loading buffer [60 mM Tris-HCl pH 6.8, 1% (w/v) SDS, 5% (v/v) glycerol, 0.005% (w/v) bromophenol blue and 1% (v/v) 2-mercaptoethanol]. Once separated by SDS-PAGE, proteins were transferred to a polyvinylidene difluoride membrane (PVDF, Immobilon-P, Merck-Millipore), then blocked and incubated with antibodies, as described [11, 12]. Antibodies and dilutions used for western blotting are described in Table S4. Membranes were developed using the Western ECL Substrate kit (Bio-Rad). The developed membranes were exposed to X-ray film or imaged with the Chemidoc Touch Imaging System (Bio-Rad).

***In vitro* mammalian cell culture.** Cell lines were grown in monolayer in culture flasks using appropriate cell culture media: HeLa cells (ATCC® CCL-2™), NIH-3T3 and Her14 mouse embryo fibroblasts [13, 14] were cultivated in Dulbecco's modified Eagle's medium (DMEM, Sigma-Aldrich), supplemented with 10% fetal bovine serum (FBS, Gibco-Thermo Fisher Scientific) (complete DMEM). HCT116 colorectal cancer epithelial cells (ATCC® CCL-247™) were grown in McCoy's 5A (Sigma-Aldrich) medium supplemented with 10% FBS (complete McCoy's). All cell lines were cultivated at 37°C with 5% $CO_2$ under static conditions. For *in vitro* cell assays, the cells were harvested from flasks at 100% confluency and placed in sterile and tissue culture-treated 6, 24 or 96-well plates (Thermo Fisher Scientific), at the desired cell concentration (~$10^5$ cells/mL). Cell lines were routinely checked for the absence of Mycoplasma spp. contamination using the Mycoplasma Gel Detection Kit (Biotools).

***In vitro* cell culture infections.** When preparing SIEC cultures for infection of mammalian cells, two experimental settings were used in relation to the induction of the T3SS components: preinduction prior to infection or induction after infection. When the T3SS was preinduced, SIEC strains were grown overnight (O/N) from a single colony in 5 mL of LB in capped Falcon tubes (BD Biosciences) at 160 rpm and 37ºC. The next day, bacterial cultures were diluted 1:100 in 5 mL of LB with the appropriate inducer (IPTG at 0.1 mM for SIEC, or aTc at 200 ng/mL for SIEC-X). Cultures were induced for 2 hours with agitation (160 rpm) at 37ºC when the cargo was induced with L-ara at 0.4%

(w/v) or with aTc at 200 ng/mL, as indicated. One hour later, cultures were diluted in serum-free DMEM containing inducers (IPTG or aTc combined with L-ara, or aTc alone) to the desired concentration and added to a monolayer of mammalian cells at ~80% confluency. Alternatively, the T3SS expression was induced over infected cells. In this case, bacteria were grown O/N from a single colony in glass flasks in 10 mL of LB under static conditions at 37ºC. Then, O/N cultures were directly diluted in serum-free DMEM containing IPTG or aTc, alone or combined with L-ara, and added over the cells. For experiments allowing bacterial adhesion prior to T3SS induction, inducers were added at the indicated time point post infection (30 or 60 minutes).

**Cytoplasmic protein extracts from infected cells.** To analyze the translocated protein in the cytoplasm of infected mammalian cells, infection with SIEC strains was carried out over mammalian cells grown in 6-well plates (Thermo Fisher Scientific). The cytoplasmic content of the infected cells was separated as follows: First, cells were extensively washed (10 times) with PBS (phosphate-buffered saline: 8 mM $Na_2HPO_4$, 1.5 mM $KH_2PO_4$, 3 mM KCl, 137 mM NaCl pH 7.0) to eliminate as many bacteria as possible. Subsequently, the dried plate containing the cells was frozen at -80ºC. On the next day, 0.5 mL of 1x lysis buffer [6.25 mM NaPPi, 1 mM NaF, 1 mM $Na_3VO_4$, 50 mM, 150 mM NaCl, Tris-HCl pH 7.4, 2 mM EDTA, 1% (v/v) NP-40 and Protease inhibitors (Sigma-Aldrich #11697498001)] was added to each well. Then, cells were scrapped and transferred to 1.5 mL microcentrifuge tubes. After 30 minutes at 4ºC, tubes were centrifuged at 21,420 *g* for 15 minutes at 4ºC. The supernatant of each tube was collected and boiled for 10 minutes with reducing SDS-PAGE loading buffer for analysis by western blot.

**β-lactamase reporter translocation assay.** In infected mammalian cells with SIEC strains, protein translocation quantification was assessed using the LiveBLAzer™ FRET B/G Loading Kit with CCF2-AM (Thermo Fisher Scientific). Mammalian cells were seeded two days prior to the experiment in a tissue culture-treated 96-well black plate with a flat clear bottom (Corning®) at a concentration of ~$10^4$ cells per well. Wells with no cells were also established as blanks. The plate was incubated at 37°C with 5% $CO_2$ under static conditions for 48 hours. Bacterial cultures were prepared to infect the cells as described above and diluted in serum-free DMEM (see *in vitro* infections). Prior to the addition of the bacteria, the mammalian cell culture was washed 3 times with serum-free DMEM to eliminate FBS from the medium. After the infection and translocation process occurred for the indicated time (usually 3 hours), wells were washed 3 times with preheated phenol red-free HBSS (Hanks Balanced Salts solution, Gibco-Thermo Fisher

Scientific). Then, the β-lactamase substrate CCF2/AM (Thermo Fisher Scientific) was added to each well at a final concentration 1 μM. Following the supplier guidelines to load cells with substrate, the plate was incubated at room temperature in the dark for 90 minutes before reading in a SpectraMax iD5 Multi-mode Microplate Reader (Molecular Devices®). The reading configuration used, following the supplier's indications, was a fluorescent dual-readout: excitation at 409 nm wavelength, emission at 450 nm (blue fluorescence) firstly, and emission at 520 nm (green fluorescence) secondly. For representation of the results, relative translocation levels were obtained by calculating the fluorescence emission ratio between 450 nm and 520 nm (blue emission/green emission) for each well. Each experimental condition was normalized by the emission ratio of the non-infected cells.

Protein translocation at the experimental endpoint was also visualized with fluorescence microscopy using a Leica TCS SP8 multispectral confocal system (Advanced Light Microscopy Service of CNB). For this purpose, IBIDI 4- or 8-well chambers were employed instead of 96-well plates. The infection process was performed as described, and in this case 0.5 mL of β-lactamase substrate mix at 1 μM was added to cover the bottom of the IBIDI® chamber. After incubating for 90 minutes in the dark, images were acquired with the confocal microscope. Both color images were taken simultaneously for each well and superimposed to visualize the final composite. Acquisition conditions were maintained for all wells. ImageJ software (NIH) was used for image composition. If time-lapse images were captured, the mammalian cells were preloaded with CCF2-AM β-lactamase substrate prior to the infection. Preloading process was adapted from [15]. Briefly, cells were cultivated in IBIDI® chambers as described above, but prior to the addition of bacteria, cells were covered with β-lactamase reagent solution containing probenecid (Sigma®) at 2.5 mM final concentration. Cells were then incubated for 60 minutes in the dark at room temperature, and subsequently for 15 minutes at 37ºC. Bacteria were then added in serum-free DMEM containing probenecid at 2.5 mM. Images were captured at different time points using a Leica TCS SP8 multispectral confocal system keeping the infected cell culture at 37ºC. When washes were performed, the fresh medium added also contained probenecid at 2.5 mM. Acquisition conditions were maintained in all wells. ImageJ software was used for time lapse composition.

**Flow cytometry of bacteria.** Bacteria corresponding to 1 $OD_{600}$ unit (ca. $3x10^8$ CFU) were centrifuged at 4,000 *g* for 3 minutes, washed twice in prefiltered PBS and finally resuspended in 1 mL of PBS containing 10% (v/v) of goat serum (Sigma). Two hundred microliters of each sample were transferred to a new tube and incubated with mouse monoclonal anti-myc antibody 9B11 (Cell Signaling Technology, diluted 1:200) for 1 hour (Table S4). Subsequently, the samples were washed three times in prefiltered PBS, resuspended in 500 µL of PBS with 10% (v/v) of goat serum and goat polyclonal Alexa Fluor 488-conjugated anti-mouse IgG antibodies (Thermo Fisher Scientific, 1:500 dilution) and incubated for 45 minutes in the dark. Finally, samples were washed three times, resuspended in 1 mL of PBS and transferred to cytometry tubes for analysis in the Flow Cytometry Service of the CNB with a Gallios Cytometer (Beckman Coulter). Results were analyzed using FlowJo software or Kaluza software (Beckman Coulter).

**Flow cytometry of mammalian cells.** Tumor cell cultures from the indicated cell lines were washed with DMEM (37ºC) and trypsinized with 1 mL of 0.05% trypsin/EDTA (Gibco) for 5 minutes. Next, the trypsin was neutralized with DMEM 10% FBS and the cells were washed twice with 1 mL of HBSS at 135 *g* for 3 minutes and resuspended in 500 µL of filtered PBS containing the primary antibody anti-EGFR (1 µg/mL, Calbiochem) and incubated for 1 hour on ice. Cells were then washed twice with 1 mL of filtered PBS and incubated with the secondary antibody goat anti-mouse Alexa 488 (1:500) in 500 µL volume for a further hour. Cells were washed twice and resuspended in 300 µL of filtered PBS for analysis in a Gallios Cytometer (Beckman Coulter). Results were analyzed using FlowJo software or Kaluza software (Beckman Coulter). Flow cytometry of cells for quantification of Live/Dead assay is described in the corresponding section.

**Bacterial adhesion assay for confocal microscopy.** Bacterial adhesion assays were adapted from [9]. Bacterial strains were cultivated O/N at 37ºC in 10 mL of LB in 50 mL glass flasks under static conditions. On the next day, the culture was pelleted at 4000 *g* for 3 minutes, washed with PBS and diluted in serum-free DMEM to a final concentration of 0.1 $OD_{600}$/mL (ca. $3x10^7$ CFU/mL). The sample was further diluted if necessary, or directly used to infect cells. Mammalian cells were seeded the day before the experiment at 35% confluence on sterile coverslips (13 mm diameter, VWR International®) placed at the bottom of the wells of a 24-well plate. The infected cell culture was incubated for 1 hour at 37ºC, 5% $CO_2$ to allow the specific attachment of the bacteria, followed by 5–10 PBS washes (1 mL per well) until no unattached bacteria was observed in the medium. Coverslips were then fixed with 4% paraformaldehyde diluted in PBS (0.5 mL per well) for 20 minutes at room temperature. After that, the paraformaldehyde was removed, and coverslips were washed 3 times with PBS. Next, coverslips were treated for 1 hour at room temperature in a wet chamber with 50 µL of PBS-10% goat serum solution containing the primary antibody for desired staining (Table S4). The coverslips were washed by immersion 15 times in a large volume of PBS (100 mL), placed again in the wet chamber and incubated for 45 minutes at room temperature with 50 μL of PBS-10% goat serum solution, with the corresponding conjugated secondary antibody. Finally, the coverslips were washed with PBS as above, the excess of liquid was removed, and the preparation was mounted with 2 μL of Prolong (Thermo Fisher Scientific) on glass slides. The samples were examined by confocal microscopy in a Leica TCS SP8 multispectral confocal system. ImageJ software was used for image composition.

**Bacterial adhesion and protein translocation assays.** This protocol was used for all the experiments where bacterial adhesion was performed prior to injection. In these experiments, bacteria adhered to target cells first, and injection of proteins by the attached bacteria is induced later. Bacteria were grown O/N from a single colony in 10 mL of LB at 37ºC under static conditions, and the next day the cultures were directly diluted in serum-free DMEM to the desired concentration and used to infect the mammalian cell cultures. After incubation for 1 hour at 37°C with 5% $CO_2$ under static conditions, unbound bacteria were removed (if indicated) by washing 5–10 times with serum-free DMEM. Finally, protein translocation by bacteria was induced with serum-free DMEM containing the inducer molecule(s) (L-ara and/or aTc, as indicated) for 4 hours. Injection levels were assessed by the β-lactamase activity assay (see above) or the culture of infected cells was continued for the indicated time (24 or 48 hours) adding gentamicin (see below).

**Live/Dead *in vitro* cell assays**. To assess *in vitro* cell death after translocation of toxins, the cell cultures were monitored at various time points up to 48 hours post infection. To prevent cell toxicity related to the excess bacterial growth during long-term experiments, bacterial infection was stopped 4 hours after induction using gentamicin. For that purpose, cultures were washed three times with DMEM, and gentamicin was first added at a high concentration (100 μg/mL) in DMEM. The plate was incubated for 1 hour at 37°C with 5% $CO_2$ under static conditions and washed with DMEM. Finally, gentamicin was added at a low concentration (10 μg/mL) diluted in complete DMEM (containing FBS 10%) and incubated at 37°C with 5% $CO_2$ under static conditions until the plate was processed.

Cell viability was assessed using the LIVE/DEAD™ Viability/Cytotoxicity Kit for mammalian cells (Thermo Fisher Scientific). If cell viability was visualized by fluorescence microscopy, reagents were added directly over the culture following the supplier indications, incubated 10 minutes in the dark at room temperature and observed with a total internal reflection fluorescence microscope (Leica DMi8 S epifluorescence system) using GFP (green, Ex: ~450nm, Em: ~510nm) and propidium iodide (red. Ex: ~528nm, Em: ~617 nm) filters. ImageJ software was used for image composition. If viability was assessed by flow cytometry, culture supernatants were saved separately and cells were harvested from the plate incubating with 0.5 mL of trypsin for 15 minutes. Trypsin neutralization was performed with the saved supernatants for each sample, followed by centrifugation at 95 *g* for 5 minutes and then staining with LIVE/DEAD reagents for 10 minutes in the dark at room temperature. Samples were then washed with PBS, resuspended in a minimum of 250 µL of PBS and transferred to cytometry tubes for analysis in the Flow Cytometry Service of the CNB in a Gallios Cytometer (Beckman Coulter) using 488 nm excitation and measuring green fluorescence emission for calcein (i.e., 530/30 bandpass) and red fluorescence emission for ethidium homodimer-1 (i.e., 610/20 bandpass). Results were analyzed using FlowJo software.

**Subcutaneous tumor model and *in vivo* studies.** Experiments with mice performed in the CNB-CSIC Animal House Facility (ref. ES280790000182) followed the protocols approved by the Ethics Committee for Animal Experimentation of CSIC and authorized by the Division of Animal Protection of the Comunidad de Madrid (project reference PROEX 074/18). Animals were handled in strict accordance with the guidelines of the European Community 86/609/CEE. The experiments performed at Synlogic were reviewed and approved by Mispro's Institutional Animal Care and Use Committee (Mispro Biotech Services, 400 Technology Square, Cambridge, MA, 02139), in compliance with Animal Welfare Act and USA and EU legislation for experimental animals procedures. Mice used in this work were athymic female Hsd:Athymic Nude-*Foxn1*$^{nu}$ mice (Envigo). Implantation of tumors was carried out by subcutaneous injection of 0.1 mL solution of ~3x10$^{6}$ HCT116 cells in PBS containing 20% (v/v) Matrigel (BD Biosciences) in the right flank of each mouse. Tumor size was monitored at different time points using calipers, and tumor volume calculated using the formula (in mm): width$^{2}$×length×0.52 until the end of the experiment. Bacteria were administered when the number of mice necessary for the experiment showed a tumor size mean of ~120 mm$^{3}$, with 90% of the mice bearing tumors between 80 mm$^{3}$ and 150 mm$^{3}$. Bacteria were grown overnight in glass flasks at 37ºC under static conditions, and on the next morning cells were washed 3 times in PBS and diluted to the indicated concentration in preheated PBS to prepare the injectable solution. To assure the concentration of this injectable solution, bacteria were counted on a Cellometer cell counter (Nextcellom Bioscience) and/or plated for next day colony counting. At day 0, bacteria were dosed by intratumoral injection of 40 μL of the bacterial suspension per mouse. This process was repeated twice (at days 3 and 6), three doses in total. Protein translocation by SIEC strains was induced by daily intraperitoneal injection of 10 μg of aTc in 0.1 mL of PBS per mouse (0.1 mL of a 100 μg/mL solution). If bacteria were administered the same day, the aTc injection was performed a minimum of 4 hours later. For determination of bacterial CFU per gram (CFU/g) of tumors, animals were euthanized and tumors were excised, placed individually into (pre-weighed) sterile tubes containing 5 mL of PBS and weighed. Samples were then transferred to sterile sampling bags (VWR) and Triton X-100 (Sigma) was added to a final concentration of 0.2% (v/v). Samples were homogenized by soft mechanical squeezing. Next, a 100 μL sample of the homogenates was serially diluted in LB, plated on LB agar and incubated overnight at 37°C to determine CFUs. Bacterial titers were expressed as CFU/g of tumor.

### Table S1. Bacterial strains.

| Name | Genotype | Reference |
|---|---|---|
| DH10B-T1R | (F- λ-) *mcrA Δmrr-hsdRMS-mcrBC φ80lacZDM15 ΔlacX74 recA1 endA1 araD139 Δ(ara, leu)7697 galU galK rpsL* (StrR) *nupG tonA* | [4], ThermoFisher |
| BW25141 | (F- λ-) *Δ(araD-araB)567, ΔlacZ4787(::rrnB-3), Δ(phoB-phoR)580, galU95, ΔuidA3::pir, recA1, endA9(del-ins)::FRT, rph-1, Δ(rhaD-rhaB)568, hsdR51* | [6] |
| MG1655 | K-12 (F- λ-) | [16] |
| EcM1 (AAEC0072) | MG1655 *ΔfimA-H rpsL*(StrR) | [17, 18] |
| SIEC | EcM1 *yfaL::*$P_{tac}$*LEE1, yeeJ::*$P_{tac}$*LEE2, yra::*$P_{tac}$*LEE3, yfc::*$P_{tac}$*escD, yebT::*$P_{tac}$*LEE4* | [10] |
| SIEC ΔescN | SIEC Δ*escN* | This work |
| SIEC Δara | SIEC Δ*araCBAD* | |
| SIEC ypjA::bla | SIECΔara *ypjA::araC*-$P_{BAD}$*_bla* | This work |
| SIECΔescN ypjA::Bla | SIEC Δara Δ*escN ypjA::araC*-$P_{BAD}$*_bla* | This work |
| SIEC ypjA::Tir-Bla | SIEC Δara *ypjA::araC*-$P_{BAD}$*_tir-bla_cesT* | This work |
| SIECΔescN ypjA::Tir-Bla | SIEC Δara Δ*escN ypjA::araC*-$P_{BAD}$*_tir-bla _cesT* | This work |
| SIEC ypjA::NleC-Bla | SIEC Δara *ypjA::araC*-$P_{BAD}$*_nleC-bla_cesT* | This work |
| SIECΔescN ypjA::NleC-Bla | SIEC Δara Δ*escN ypjA::araC*-$P_{BAD}$*_nleC-bla_cesT* | This work |
| SIEC ypjA::EspH-Bla | SIEC Δara *ypjA::araC*-$P_{BAD}$*_espH-bla_cesT* | This work |
| SIECΔescN ypjA::EspH-Bla | SIEC Δara Δ*escN ypjA::araC*-$P_{BAD}$*_espH-bla_cesT* | This work |
| SIEC ypjA::Map-Bla | SIEC Δara *ypjA::araC*-$P_{BAD}$*_map-bla_cesT* | This work |
| SIECΔescN ypjA::Map-Bla | SIEC Δara Δ*escN ypjA::araC*-$P_{BAD}$*_map-bla_cesT* | This work |
| SIEC ypjA::Tir30-Nb-Bla | SIEC Δara *ypjA::araC*-$P_{BAD}$*_tir30-nbGFP-bla* | This work |
| SIECΔescN ypjA::Tir30-Nb-Bla | SIEC Δara Δ*escN ypjA::araC*-$P_{BAD}$*_tir30-nbGFP-bla* | This work |
| SIEC ypjA::Tir100-Nb-Bla CesT | SIEC Δara *ypjA::araC*-$P_{BAD}$*_tir100-nbGFP-bla_cesT* | This work |
| SIECΔescN ypjA::Tir100-Nb-Bla CesT | SIEC Δara Δ*escN ypjA::araC*-$P_{BAD}$*_tir100-nbGFP-bla_cesT* | This work |
| SIEC ypjA::Tir100-NbGFP-Bla | SIEC Δara *ypjA::araC*-$P_{BAD}$*_tir100-nbGFP-bla* | This work |
| EcM1*lux* SAegfr | EcM1 *matB::P2-luxCDABE flu::*$P_{N25}$*-SAegfr* | This work |
| EcM1*lux* SAtir | EcM1 *matB::P2-luxCDABE flu::*$P_{N25}$*-SAtir* | This work |
| SIEC-X | SIEC Δara ΔlacI *csgCG::3R-X* (*tetR-Ptet-*$cI^{+}Ind^{-}$, $P_R$-$lacI^{W220F-ASV}$) | [12] |
| SIEC-X SAtir | SIEC-X *flu::*$P_{N25}$*-SAtir* | This work |
| SIEC-X SAegfr | SIEC-X *flu::*$P_{N25}$*-SAegfr* | This work |
| SIEC-X SAegfr ΔescN | SIEC-X SAegfr Δ*escN* | This work |
| SIEC-X SAegfr Tir100-Nb-Bla | SIEC-X SAegfr *ypjA::araC*-$P_{BAD}$*_tir100-nbGFP-bla_cesT* | This work |
| SIEC-X SAegfr Tir100-PE25-Bla | SIEC-X SAegfr *ypjA::araC*-$P_{BAD}$*_tir100-PE25-bla_cesT* | This work |
| SIEC-X SAegfr ΔescN Tir100-PE25-Bla | SIEC-X SAegfr Δ*escN ypjA::araC*-$P_{BAD}$*_tir100-PE25-bla_cesT* | This work |
| SIEC-X SAegfr Tir100-TccC3-Bla | SIEC-X SAegfr *ypjA::araC*-$P_{BAD}$*_tir100-*$tccC3_{hvr}$*-bla_cesT* | This work |
| SIEC-X SAegfr ΔescN Tir100-TccC3-Bla | SIEC-X SAegfr ΔescN *ypjA::araC*-$P_{BAD}$ *tir100- *$tccC3_{hvr}$*-bla_cesT* | This work |
| SIEC-X SAegfr Ptet-Tir100-Nb | SIEC-X SAegfr *ypjA::*$P_{tet}$*_tir100-nbGFP_cesT* | This work |
| SIEC-X SAegfr Ptet-Tir100-PE25 | SIEC-X SAegfr *ypjA::*$P_{tet}$*_tir100-PE25_cesT* | This work |
| SIEC-X SAegfr Ptet-Tir100-TccC3 | SIEC-X SAegfr *ypjA::*$P_{tet}$*_tir100-*$tccC3_{hvr}$*_cesT* | This work |
| SIEC-X SAegfr ΔescN Ptet-Tir100-PE25 | SIEC-X SAegfr Δ*escN ypjA::Ptet_tir100-PE25_cesT* | This work |
| SIEC-X SAegfr ΔescN Ptet-Tir100-TccC3 | SIEC-X SAegfr Δ*escN ypjA::Ptet_tir100-*$tccC3_{hvr}$*_cesT* | This work |
| SIEC-X SAtir Ptet-Tir100-Nb | SIEC-X SAtir *ypjA::Ptet_tir100-nbGFP_cesT* | This work |
| SIEC-X SAtir Ptet-Tir100-PE25 | SIEC-X SAtir *ypjA:: Ptet _tir100-PE25_cesT* | This work |
| SIEC-X SAtir Ptet-Tir100-TccC3 | SIEC-X SAtir *ypjA:: Ptet _tir100-*$tccC3_{hvr}$*_cesT* | This work |

**Table S2. Plasmids.**

| Name | Features / use | Reference |
|---|---|---|
| pACBSR-Sp | $Sp^R$, p15A ori, *araC*, $P_{BAD}$, *I-SceI* and *λ Red* genes | 10 |
| pGE | $Km^R$; R6K ori (suicide vector) | 9 |
| pGEΔescN | pGE to delete *escN* | 19 |
| pGEΔara | pGE to delete *araCBAD* operon in MG1655 | This work |
| pGEypjA | pGE with HRs flanking ypjA of MG1655 | This work |
| pGEypjA_$P_{BAD}$-*bla* | pGEypjA for integration of *araC*-$P_{BAD}$-*bla* | This work |
| pGEypjA_*araC*-$P_{BAD}$_Tir-Bla_CesT | pGEypjA for integration of *araC*-$P_{BAD}$_tir-bla_cesT | This work |
| pGEypjA_*araC*-$P_{BAD}$_NleC-Bla_CesT | pGEypjA for integration of *araC*-$P_{BAD}$_NleC-bla_cesT | This work |
| pGEypjA_*araC*-$P_{BAD}$_EspH-Bla_CesT | pGEypjA for integration of *araC*-$P_{BAD}$_espH-bla_cesT | This work |
| pGEypjA_*araC*-$P_{BAD}$_MAP-Bla_CesT | pGEypjA for integration of *araC*-$P_{BAD}$_MAP-bla_cesT | This work |
| pBAD18-Kan | $Km^R$, pBR322 ori, *araC*-$P_{BAD}$ | 20 |
| pBAD_Bla | pBAD18-Kan, *araC*-$P_{BAD}$_bla | This work |
| pBAD_Tir30-Nb-Bla | pBAD18-Kan, *araC*-$P_{BAD}$_*tir30-Nb-bla* | This work |
| pBAD_Tir100-Nb-Bla_CesT | pBAD18-Kan, *araC*-$P_{BAD}$_tir100-Nb-bla_cesT | This work |
| pBAD_EspF20-Bla | pBAD18-Kan, *araC*-$P_{BAD}$_espF20-bla | This work |
| pBAD_EspF20-Nb-Bla | pBAD18-Kan, *araC*-$P_{BAD}$_espF20-nb-bla | This work |
| pGEypjA_*araC*-$P_{BAD}$_Tir30-Nb-Bla | pGEypjA for integration of *araC*-$P_{BAD}$_tir30-nb-Bla | This work |
| pGEypjA_*araC*-$P_{BAD}$-Tir100-Nb-Bla_CesT | pGEypjA for integration of *araC*-$P_{BAD}$_tir100-nb-Bla_cesT | This work |
| pGEypjA_*araC*-$P_{BAD}$-Tir100-Nb-Bla | pGEypjA for integration of *araC*-$P_{BAD}$_tir100-nb-bla | This work |
| pBAD_*araC*-$P_{BAD}$_Tir100-BIM-Bla_CesT | pBAD18-Kan, *araC*-$P_{BAD}$_tir100-BIM-bla_cesT | This work |
| pBAD_*araC*-$P_{BAD}$_Tir100-tBID-Bla_CesT | pBAD18-Kan, *araC*-$P_{BAD}$_Tir100-tBID-bla_cesT | This work |
| pBAD_*araC*-$P_{BAD}$_Tir100- GrzB-Bla_CesT | pBAD18-Kan, *araC*-$P_{BAD}$_tir100-granzymeB-Bla_cesT | This work |
| pBAD_*araC*-$P_{BAD}$_Tir100-OVA-Bla_CesT | pBAD18-Kan, *araC*-$P_{BAD}$_tir100-OVA-bla_cesT | This work |
| pBAD_*araC*-$P_{BAD}$_Tir100-Survivin-Bla_CesT | *araC*-$P_{BAD}$_tir100-survivin-bla_cesT | This work |
| pBAD_*araC*-$P_{BAD}$_Tir100-TK-Bla_CesT | pBAD18-Kan, *araC*-$P_{BAD}$_tir100-TK-bla_cesT | This work |
| pBAD_*araC*-$P_{BAD}$_Tir100-Sox2-Bla_CesT | pBAD18-Kan, *araC*-$P_{BAD}$_tir100-sox2-bla_cesT | This work |
| pGEcsg3R-X | pGE*csg* for integration of 3R-X | 12 |
| pGErecomb_SAegfr2 | pGE with flanking HRs 3'*eae* and 3'*flu* for insertion of Vegfr2 in SA | This work |
| pBAD_Tir100-Nb_CesT | pBAD18-Kan, *araC* $P_{BAD}$ tir100-nbGFP cesT | This work |
| pGEypjA_*araC*-$P_{BAD}$_Tir100-PE25-Bla_CesT | pGE*ypjA* for integration of *araC*-$P_{BAD}$_tir100-PE25-bla_cesT | This work |
| pGEypjA_*araC*-$P_{BAD}$_Tir100-TccC3hvr-Bla_CesT | pGE*ypjA* for integration of *araC*-$P_{BAD}$_tir100-TccC3hvr-bla_cesT | This work |
| pGEypjA_Ptet_Tir100-Nb-Bla_CesT | pGE*ypjA* for integration of Ptet_tir100-Nb-bla_cesT | This work |
| pGEypjA_Ptet_Tir100-PE25_CesT | pGE*ypjA* for integration of Ptet_tir100-PE25_cesT | This work |
| pGEypjA_Ptet_Tir100-TccC3hvr_CesT | pGE*ypjA* for integration of Ptet_tir100-TccC3hvr_cesT | This work |

### Table S3. Oligonucleotides.

| Nº | Name | Sequence (5'->3') | Use |
|---|---|---|---|
| 1 | F_XbaI_RBS_MAP | TCGAGTCTAGAAAGAAGGAGATATACATATGTTTAGTCCAACGGCAATGGTAGGTAG | Cloning MAP effector |
| 2 | R_NotI_MAP | CCGATGCGGCCGCCAGCCGAGTATCCTGCACATTGTC | Cloning MAP effector |
| 3 | ypjA_5' | AGCTGTGCGAACGTGGTATTAACTTTC | Anneals with 5' HR of ypjA, for checking integrations |
| 4 | ypjA_3' | AACAACACTATGGCCTGACACTGAACG | Anneals with 3' HR of ypjA for checking integrations |
| 5 | R_AraC_test | CGATCAACTCTATTTCTCGCGGGTA | Anneals with araC for checking integrations |
| 6 | F_CesT_test | GTGGGCCATACCTGTGTTATGAGTCAG | Anneals within cesT, used for integration checking |
| 7 | F_SacI_RBS_CesT | AATTCGAGCTCAAGAAGGAGATATACATATGTCATCAAGATCCGAAC | Cloning cesT |
| 8 | F_SfiI_GranzymeB | ATTGCGGCCCAGCCGGCCATGATTATTGGCGGCCATGAAGC | Cloning GranzymeB |
| 9 | R_NotI_GranzymeB | CCGATGCGGCCGCATAGCGTTTCATGGTTTTTTTAATCCAATG | Cloning GranzymeB |
| 10 | F_SfiI_OVA | ATTGCGGCCCAGCCGGCCATGGGCAGCATTGGCGCG | Cloning OVA |
| 11 | R_NotI_OVA | CCGATGCGGCCGCCGGGCTCACGCAGCGG | Cloning OVA |
| 12 | F_SfiI_TK | ATTGCGGCCCAGCCGGCCATGGCTTCGTACCCCTGCCATC | Cloning TK |
| 13 | R_NotI_TK | CCGATGCGGCCGCGTTAGCCTCCCCCATCTCCCG | Cloning TK |
| 14 | F_SacI_RBS_EspF20 | AATTCGAGCTCAAGCTTAAGAAGGAGATATACATATGCTTAATGGAATTAGTAACGCTGCTTC | Cloning EspF20 |
| 15 | R_SpeI_Bla | CATGCACTAGTTTACCAATGCTTAATCAGTGAGGCACC | Cloning EspF20 |
| 16 | R_NotI_BIM | CCGATGCGGCCGCATGCATGCGCCACACCAGG | Cloning BIM |
| 17 | F_SfiI_BIM | GGTGCGGCCCAGCCGGCCGCGAAACAGCCGAGCGATG | Cloning BIM |
| 18 | F_SfiI_Sox2 | GGTGCGGCCCAGCCGGCCATGATGGAGACGGAGCTGAAGCC | Cloning Sox2 |
| 19 | R_NotI_Sox2 | CCGATGCGGCCGCCATGTGCGACAGGGGCAGTG | Cloning Sox2 |
| 20 | R_cargo_seq | GCGTCGAAGCATGGTGATGGT | DNA sequencing of cargoes |
| 21 | R_NotI_tBID | CCGATGCGGCCGCATCCATACCATTACGTGCCAGGCTAC | Cloning tBID |
| 22 | F_LacI_seq | GCGTCTGCGTCTGGCTGGC | For lacI sequencing |
| 23 | F_SfiI_PE25 | GGTGCGGCCCAGCCGGCCGGCCCGGCGGACAGC | Cloning PE25 |
| 24 | R_NotI_PE | CCGATGCGGCCGCCTTCAGGTCCTCGCGCGG | Cloning PE25 |
| 25 | R_BglII_STOP_PE25 | TTCTTAGATCTTCACTTCAGGTCCTCGCGCGGCG | For cloning PE25 with STOP codon |
| 26 | F_SfiI_TccC3 | GGTGCGGCCCAGCCGGCCATGCCGACCATTGCAGAACGTATT | For cloning TccC3 |
| 27 | R_BglII_STOP_TccC3 | TTCTTAGATCTTCAACGTTTATGCGGTTTCACATTTTTCA | For cloning TccC3 with STOP codon |

### Table S4. Antibodies and fluorophores.

| Use | Antigen | Primary antibody | Secondary Antibody |
|---|---|---|---|
| **Western blot** | | | |
| | **Bla** | anti-Bla mAb (QED Bioscience, 1:2000) | anti-mouse-POD (Sigma, 1:5000) |
| **Cytometry** | | | |
| | **Myc tag** | anti-Myc tag mAb (Cell Signaling Technology, 1:500) | anti-mouse-Alexa488 (Life Technologies, 1:500) |
| **Microscopy** | | | |
| | ***E. coli*** | rabbit anti-E.coli K-12 (Biodesign, 1:1000) | anti-rabbit 488 (Life Technologies, 1:500) |
| | **DNA** | DAPI (Life Technologies, 1:1000) | -- |
| | **EGFR** | anti-EGFR mouse monoclonal mAb (1:500, Calbiochem-Merck Millipore REF: GR01) | anti-mouse-Alexa (1:500, Life Technologies) |

### Supporting Figures

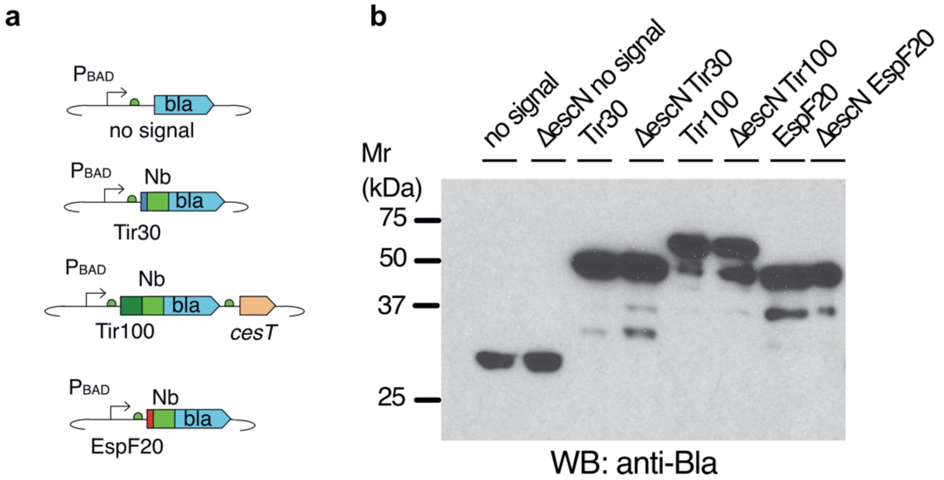


**Figure S1. Expression of Bla fusions with different T3S signals from multicopy pBAD plasmids. a,** Scheme of the Bla fusions in pBAD plasmids having different T3S signals, Tir30, Tir100, EspF20, Nb, and *cesT*, as indicated. **b,** Western blot (WB) of whole-cell protein extracts from induced (L-ara, IPTG) SIEC bacteria carrying the indicated T3S signal in a Bla fusion. WB incubated with anti-Bla ($\beta$-lactamase) antibody and secondary anti-mouse IgG-peroxidase conjugate.

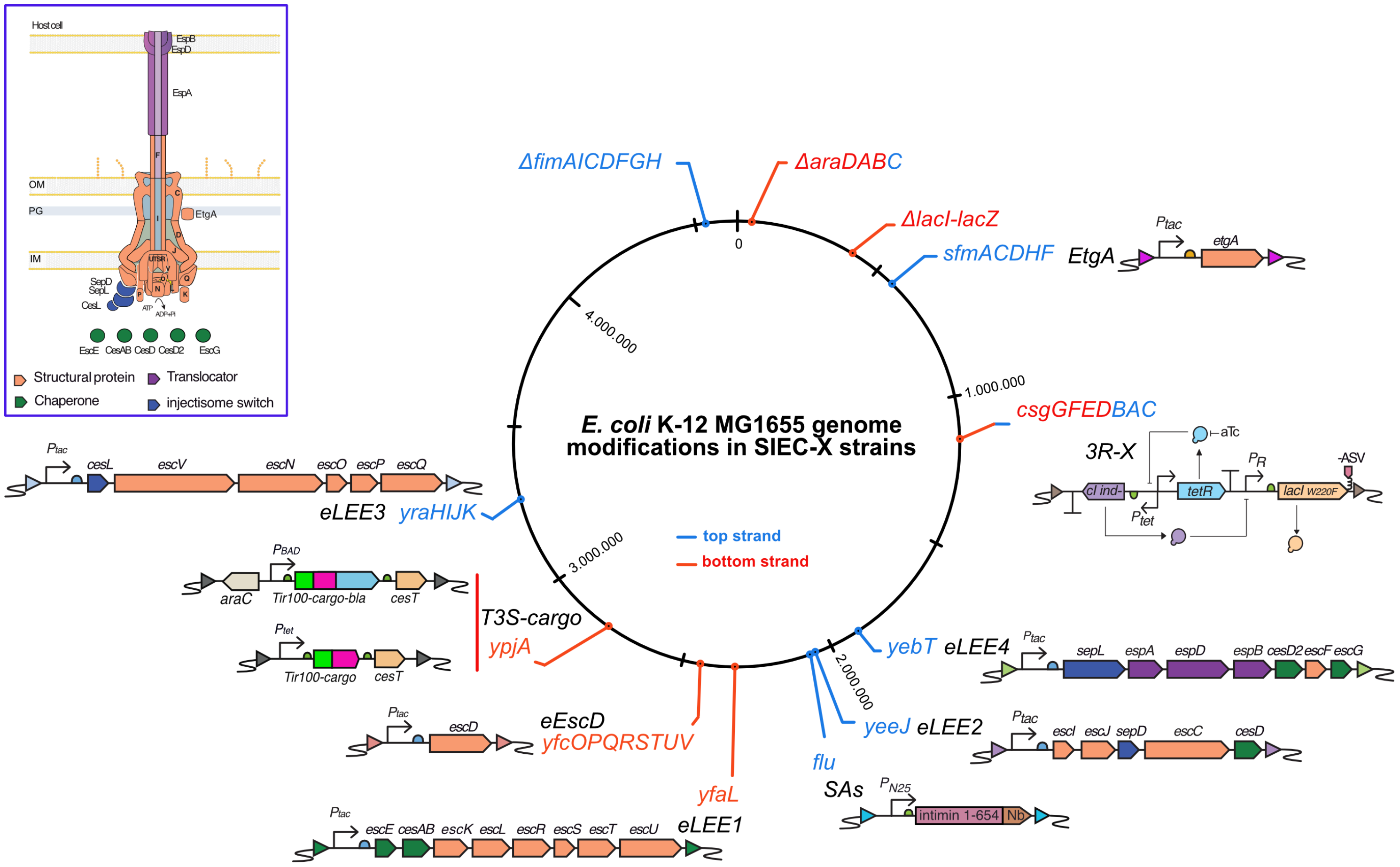


**Figure S2. Genome modifications in SIEC-X strains.** Scheme indicating markerless gene insertions and deletions found in SIEC-X strains over the circular chromosome of the reference *E. coli* K-12 MG1655 strain. Orientation of the coding regions in the top or bottom strands of DNA is labeled. Top left inset shows a scheme of the type III secretion system (T3SS) injectisome of SIEC assembled in the bacterial envelope (inner membrane, IM; peptidoglycan, PG; outer membrane, OM) for the translocation of cargo proteins from the cytoplasm of bacteria to the host cell.

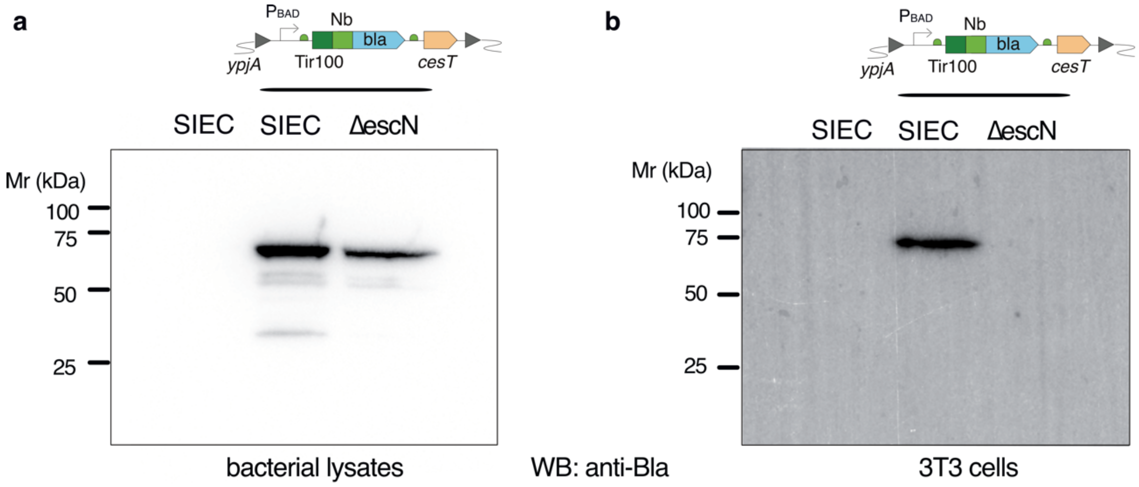


**Figure S3. Translocated Tir100-Nb-Bla fusion detected in the cytoplasm of infected 3T3 cells.** Western blot (WB) of: **a**, bacterial lysates of induced (L-ara, IPTG) parental SIEC, SIEC with integrated *ypjA*::*araC*_$P_{BAD}$_Tir100-Nb-bla_*cesT*, and isogenic Δ*escN* mutant. **b**, Cytoplasmic protein extracts of 3T3 cells after infection with bacteria from the indicated strains. **a, b,** WB incubated with anti-Bla antibody, as in Fig. S1.

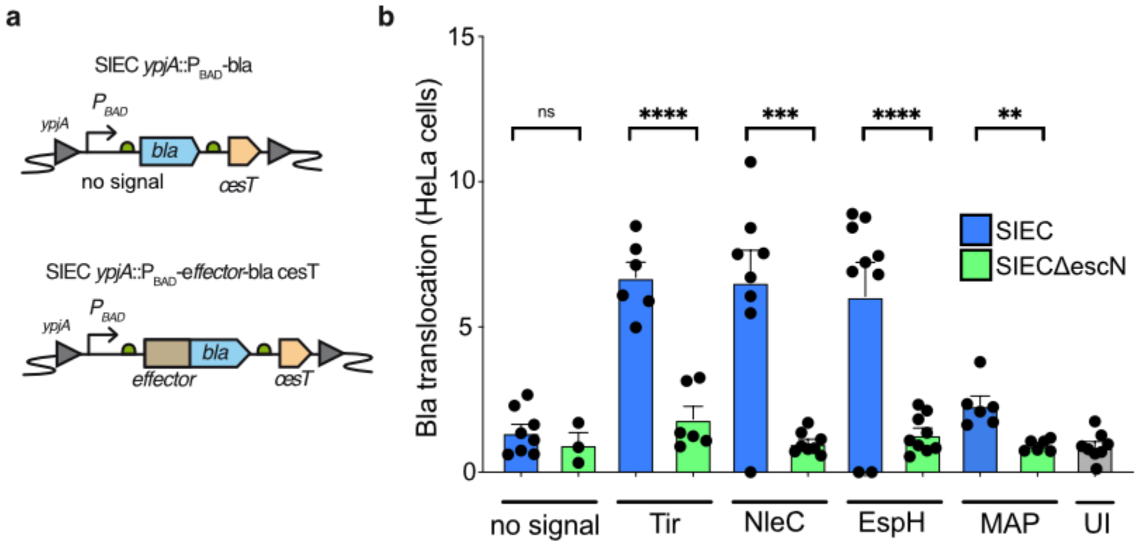


**Figure S4. Translocation of natural EPEC effectors by SIEC. a,** Schematic representation of the constructs integrated into the *ypjA* locus for the coexpression under $P_{BAD}$ regulation of natural EPEC effector-Bla fusion and CesT chaperone. **b**, Translocation levels to HeLa cells of the effectors Tir, NleC, EspH and Map after infection with induced (L-ara, IPTG) SIEC strains carrying the corresponding integrated construct. Data are presented as the mean ± s.e. Unpaired two-tailed t-tests were used to evaluate differences between groups. (*) $P<0.05$, (**) $P<0.01$, (***) $P<0.001$, (****) $P<0.0001$. Each condition was compared with its correspondent Δ*escN* control: no signal ($P=0.4383$), Tir ($P<0.0001$), NleC ($P<0.0001$), EspH ($P<0.0001$), MAP ($P=0019$).

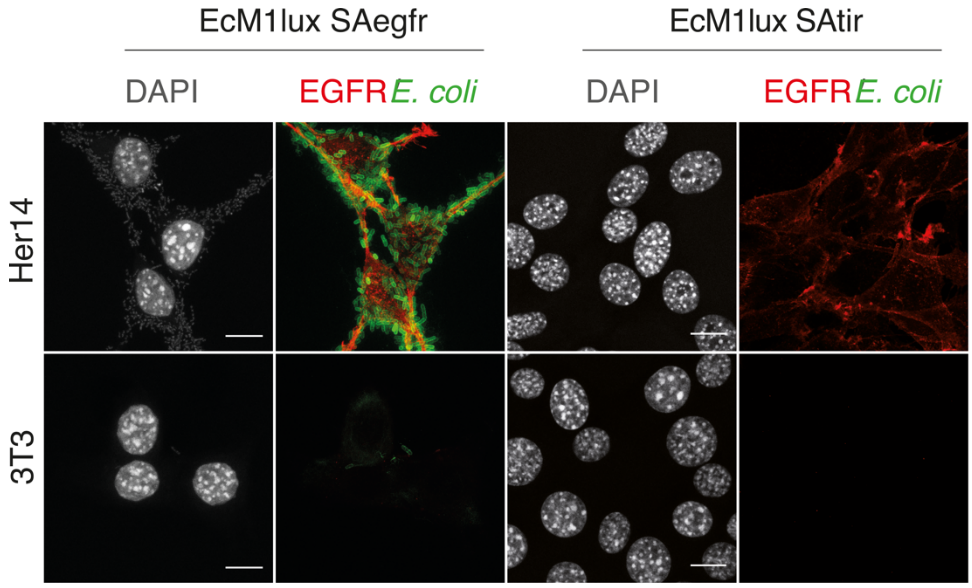


**Figure S5. Adhesion to EGFR+ cells of *E. coli* bacteria expressing SAegfr.** Confocal microscopy images of Her14 (top panel) and 3T3 cells (bottom panel) infected with EcM1*lux*SAegfr or EcM1*lux*SAtir (control), as indicated. Bacteria were stained with anti-*E. coli* polyclonal serum and anti-rabbit IgG-Alexa 488 (green), EGFR was stained with anti EGFR mAb and anti-mouse IgG-Alexa-594 (red), cell nuclei were stained with DAPI (grey). Scale bars (white lines) represent 10 μm.

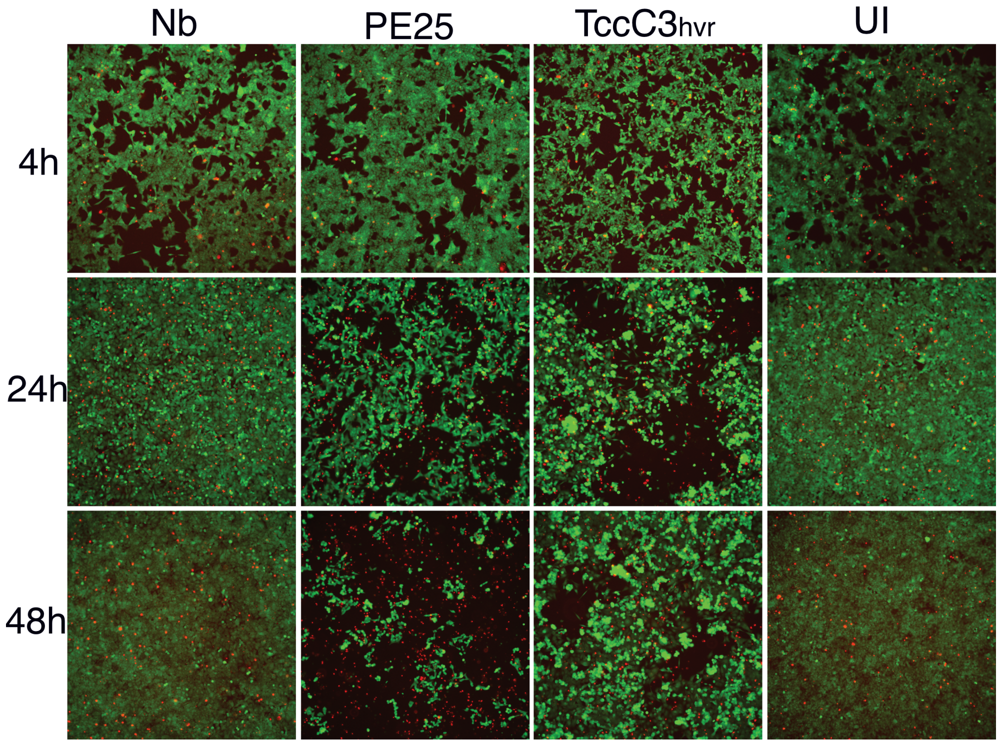


**Figure S6. Time-course of tumor cell death and cytotoxicity caused by the translocated ART toxins.** Microscopy images of HCT116 cells infected with SIEC-X SAegfr strains translocating the indicated ART toxin fragment (PE25, $TccC3_{hvr}$), or Nb as control, and stained with the Live/Dead assay (Calcein-AM/Ethidium homodimer) 4 h, 24 h, and 48 h post infection, as in Figure 3.

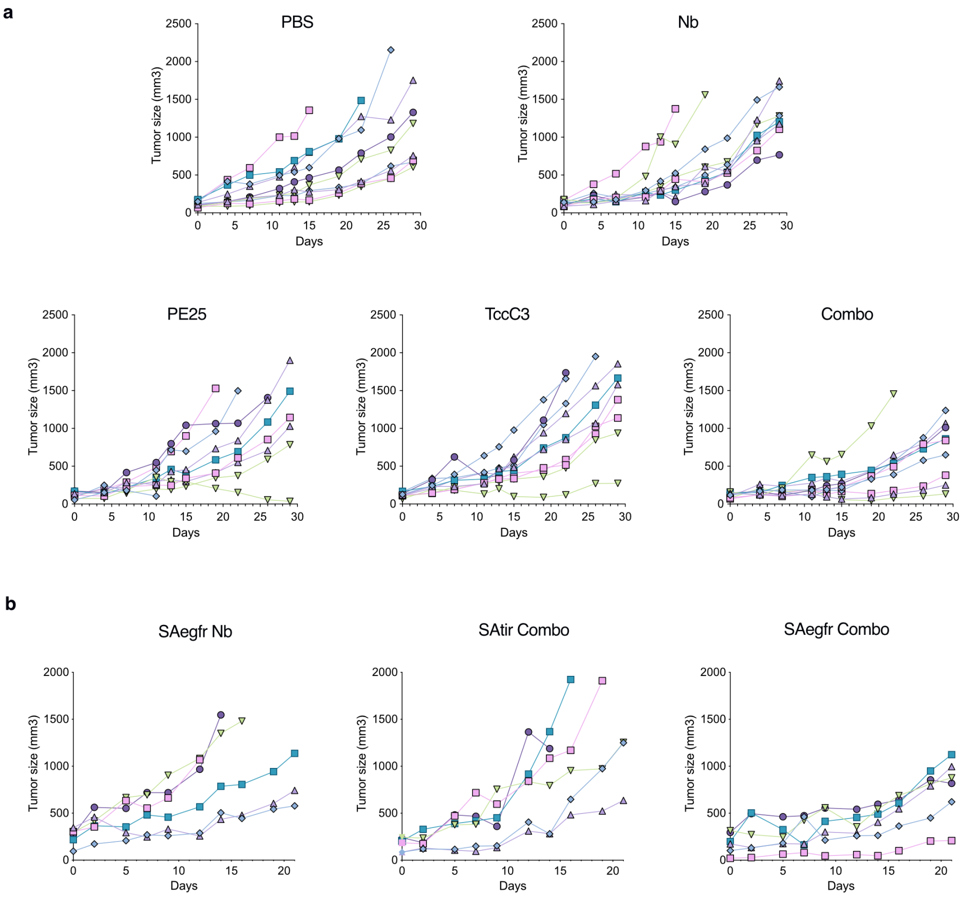


**Figure S7. Growth curves of each individual tumor in experimental groups.** Graphs show the growth of each tumor in the indicated experimental groups corresponding to *in vivo* experiments in **a**, Figure 4b,c. **b,** Figure 4d,e.

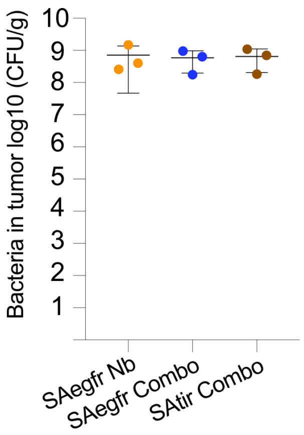


**Figure S8. Bacterial counts in tumors.** Graph shows colony forming units per gram of tumor (CFU/g) in tumors from the indicated experimental groups at the end of experiment in Fig. 4e (day 21). Counts from serial dilutions of tumor homogenates plated in LB agar plates for each experimental group. Data are presented as the mean ± s.e. Unpaired two-tailed t-tests were used to evaluate differences between groups. (*) $P$<0.05, (**) $P$<0.01, (***) $P$<0.001, (****) $P$<0.0001. SAegfr Nb versus SAegfr Combo ($P$=7986), SAegfr Nb versus SAtir Combo ($P$=6080), SAegfr Combo versus SAtir Combo ($P$=4416).
